# History-dependent ephaptic interactions in paired olfactory receptor neurons

**DOI:** 10.64898/2026.01.23.701383

**Authors:** Lydia Ellison, György Kemenes, Thomas Nowotny

## Abstract

Olfactory sensing begins with the transduction of odors into receptor currents on the dendrites of olfactory receptor neurons (ORNs). In insects and many other arthropods, ORNs are grouped stereotypically in hair-like sensilla on the surface of olfactory organs, enabling mutual inhibition through non-synaptic ‘ephaptic’ interactions (NSIs). Given the electrical, and therefore virtually instantaneous, nature of NSIs, it has been hypothesized that they contribute to processing fast temporal elements of mixed odor plumes. Here, we present single sensillum recordings and computational modeling that characterize NSIs during short offset dual-odor stimulations in the olfactory sensilla of adult female *Drosophila melanogaster*. We find in the experiments that the magnitude of inhibition between co-housed ORNs cannot be predicted by their instantaneous activity (firing rate) alone. It is adaptation-dependent, with strong effects only occurring when the inhibited ORN is adapted. This limits the usefulness of NSIs for fast odor processing when ORNs lack time to adapt. We reproduced the observed phenomena in a computational model and use this model to explain how the adaptation-dependence of NSI-mediated inhibition arises from nonlinearities in neural responses. We conclude that NSIs are unlikely to support the encoding of fast temporal dynamics in mixed odor stimuli, instead contributing to slower peripheral processing, supporting roles such as novelty detection. More broadly, we demonstrate how the nonlinear interactions of fairly simple electrical components lead to non-intuitive results, offering insight into the longstanding debate around ephaptic interactions in other systems, such as the mammalian CNS.

## Introduction

Nervous systems fundamentally exist to enable animals to generate appropriate behavior in response to environmental conditions. Accordingly, animals possess specialized neural structures for detecting relevant stimuli. In all but the simplest systems, these primary sensory structures relay information to a central nervous system (CNS), where relevant features are extracted. However, sensory information is also often pre-processed in the periphery, a phenomenon frequently overlooked. Many identified examples of peripheral sensory interaction are enabled by neural compartmentalization, where the close proximity of neurons allows cross-talk that serves diverse roles including coding stimulus identity (***Takeichi et al., 2018***), broadening the range of encoded information (***Ng et al., 2020***), enhancing valence contrast (***Wu et al., 2022; Puri et al., 2024***) and aiding in the interpretation of complex stimuli (***Stierle et al., 2013; Pannunzi and Nowotny, 2021; Bandyopadhyay and Sachse, 2023***).

In insects, non-synaptic ‘ephaptic’ interactions (NSIs) are a well-documented feature of compartmentalized olfactory receptor neurons (ORNs) (***Su et al., 2012; Zhang et al., 2019***). ORNs are housed in groups, typically of 1-4, with their highly branched dendrites extending together into sensilla – finger-like protrusions of the cuticle exoskeleton, reviewed in ***Schmidt and Benton (2020***). The restricted, electrically isolated sensillar compartment facilitates the generation of a local field potential (LFP). This LFP mediates lateral inhibition between co-housed ORNs, similar to that observed between unmyelinated neurons in the vertebrate brain (***Faber and Korn, 1989; Jefferys, 1995***), photoreceptors of compound eyes (***Shaw, 1975; Weckström and Laughlin, 2010***) and recently described between paired gustatory receptor neurons (***Lee et al., 2024, 2025***). In olfactory sensilla, the LFP is proposed to be the primary driving force for odorant-induced receptor currents (***Vermeulen and Rospars, 2004; Kaissling, 1986***). It is therefore supposed that during NSIs, LFP deflections induce lateral inhibition between ORNs by reducing the driving force acting on receptor currents (***Su et al., 2012; Zhang et al., 2019***). It is worth noting that this type of inhibition arises from reduced excitability at the dendrites due to a reduced reversal potential of the receptor current, not via a direct hyperpolarisation of the neuron.

Strong lateral inhibition between ORNs has been demonstrated across multiple sensillum types and linked to behavioral effects (***Su et al., 2012; Zhang et al., 2019; Wu et al., 2022***). In addition to the tuning of NSIs by ORN asymmetry (***Zhang et al., 2019***), the magnitude of these interactions is also modulated by stimulus-induced variations in the activity levels (firing rates) of both ORNs, with increased activity of either leading to increased NSIs (***Su et al., 2012***). The activity of the ‘inhibiting’ ORN scales the LFP deflection, while the activity of the ‘inhibited’ ORN scales membrane conductance, determining its sensitivity to LFP-mediated changes in the driving force. Although some evidence suggests that simultaneous ORN activation induces lateral inhibition (***Andersson et al., 2010; Xu et al., 2019***), detailed investigations have focused on conditions where the inhibited ORN exhibits a stable, adapted firing rate (***Su et al., 2012; Zhang et al., 2019***). The proposal that NSIs aid in processing rapidly changing mixed odors (***Vermeulen and Rospars, 2004; Pannunzi and Nowotny, 2021; Bandyopadhyay and Sachse, 2023***) rests on the assumption that NSI strength remains relevant under such dynamic conditions: during rapidly fluctuating odors, ORNs undergo pronounced adaptation-driven changes in firing rate from very high rates upon initial odor encounters, to low rates following adaptation. For NSIs to meaningfully contribute to fast temporal coding, they must scale in a way that they remain relevant for the high firing rate regimes of unadapted neurons.

In this paper, we present electrophysiology experiments with stimulation of both ORNs within a sensillum in rapid succession. These stimuli mimic the timing of overlapping odor filaments in a mixed-source plume (***Celani et al., 2014***). We thereby provide a direct test of the potential role of NSIs in processing rapidly changing odor signals. As NSIs have been documented across various sensillum types, this investigation focuses on the easily accessible ab3A, one of the large basiconic sensilla specialized for detecting food odors (***Hallem and Carlson, 2006; Gomez-Diaz et al., 2018***). Our analysis specifically targets the responses of the larger A ORN during B-to-A inhibition for two main reasons: first, the larger spike size of the A ORN reduces the risk of errors that could arise from incorrect spike sorting; and second, the inhibition of A ORNs is arguably more behaviorally important due to their role in mediating attraction, a role which is prioritized via morphological differences (***Zhang et al., 2019***) and conserved across insect species (***Cossé et al., 1998; Chang et al., 2016; Wu et al., 2022***).

## Materials and Methods

### *Drosophila* maintenance

All experiments were performed on mated adult female wild-type Oregon-Red flies, 4–8 days after eclosion. Flies were reared at 25 °C in an incubator with a 12-h light–dark cycle.

### Single-sensillum recordings

Fly antennal preparations and single-sensillum recordings (SSRs) were performed essentially as previously described (***Benton and Dahanukar, 2011; Olsson and Hansson, 2013; Pellegrino et al., 2010***). In brief, flies were immobilized with one antenna stabilized between a tapered glass microcapillary tube and a coverslip. An electrolytically sharpened tungsten electrode was inserted through the cuticle of an olfactory sensillum, bringing the tip of the electrode into contact with the sensillar lymph surrounding the ORN dendrites. The sample sizes (n) indicated in figure legends correspond to biological replicates (different sensilla), with 1-3 recorded per fly.

Electrical signals were amplified (1000X) in two stages. Firstly by connecting the recording and reference electrodes to a pre-amplifier headstage (NL100AK AC, Digitimer, Welwyn Garden City) and secondly using an AC pre-amplifier (NL104A AC, Digitimer). Signals were digitized via a 10 kHz 16-bit analogue to digital converter (Micro 1401 mk II, CED, Oxford), acquired in Spike2 software (CED, Oxford) and bandpass filtered offline (195-3000 Hz).

### Spike sorting

Automated sorting in Spike2 was not possible due to the high degree of amplitude change during high firing rates, as well as the alteration of spike shapes due to overlapping spike shapes, previously discussed in ***Ellison et al. (2025***). The first factor prevented all spikes from a single ORN being assigned to the same cluster and the second led to spike shapes that were too far outside the templates used for automated sorting. Therefore, unless otherwise stated, SSR data was predominantly sorted manually by shape in Spike2.

In brief, WaveMark channels were generated from recorded traces, identifying candidate spike events by thresholding. These markers were sorted manually by a combination of further thresholding and the close-up examination of individual spike shapes. The resulting sorted WaveMark channels corresponding to sorted A and/or B spikes were converted to Event time markers corresponding to the positive peak of each spike marker.

Responses to the simulated plume stimuli in **Figure 9** were analyzed semi-automatically using the Python-based SSSort 2.0 spike sorting algorithm (***Ellison et al., 2025***). The core algorithm was first run to detect and sort spikes with manual approval of cluster merges. The post-processing extension was then run with fully automated parameters to re-sort all spikes by considering if they could be composite spikes. Parameters were then adjusted to semi-automated post-processing mode. High firing rate regions of the data were re-analysed, with manual checks for the more difficult to sort spikes.

The accuracy of both methods has been validated in ***Ellison et al. (2025***) by the sorting of artificial data where the spike times of both neurons are known.

### Odor stimuli

Odors selective for each ORN were chosen based on previous work (***Su et al., 2012***). Chemicals were > 99% pure or of the highest purity available at Sigma-Aldrich and diluted (v/v) in mineral oil unless otherwise specified. Odor stimuli were delivered using a purpose-built high precision two-channel odor stimulator (modified from ***Raiser et al. (2017***)). To ensure stable concentrations, odor dilutions were continuously mixed during experiments and fresh dilutions made weekly. Each odor was delivered by a continuous clean air stream (250 mL/min) from a pressured tank passing into a 20 mL glass vial with a Teflon septum containing 3 mL of the diluted odorant. Each odorized air stream was then delivered to a three-way Teflon solenoid valve (LHDA1233415H, Lee Company, Westbrook, CT), which, when switched during odor stimulation, directed the odorized air into the main stream (1.5 L/min) from the same air tank via the mixing chamber of the 3D printed odor block. Two additional valved clean air streams provided balance to maintain stable air flow. These valves were controlled via a microcontroller platform (ESP32-WROOM-32, Espressif Systems, Shanghai) and custom Arduino routines. During SSRs, odor stimuli were delivered by placing the 6mm diameter glass odor delivery tube 5-10mm from the head of the immobilized fly. At this distance, PID measurements confirm that the concentration of the odor plume is stable and only minimally reduced at the maximum distance by 5% compared to the concentration measured inside the outlet tube. Opposite from the odor outlet, a slow extractor fan placed 10 cm from the fly minimized odor pulse disturbance whilst ensuring that the odor is removed and a clean background maintained. In order to ensure consistency, odor stimuli were delivered following a minimum of 100 s manual background flush with odor channel directing air to waste. At this time point most odors have reached a stable headspace concentration in the vial.

### Computational model

We simulated two Hodgkin-Huxley-type model neurons (***Traub and Miles, 1991***). In order to simulate the electrical separation between the ORN soma and dendrites, each neuron consisted of two coupled compartments. The dendritic compartments are further influenced by an additional transmembrane potential, simulating the arrangement of two ORNs co-housed in a single sensillum. The model is conceptually summarized in **Figure 4A**.

Each soma compartment, defined by its transmembrane potential *V*, was modeled using the standard Hodgkin-Huxley formalism with sodium *I*_Na_, potassium *I*_K_, and *I*_leak_ currents:

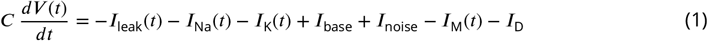

Here the leak current, simulating the flow of ions between each soma and the hemolymph through non-specific ion channels, is given by *I*_leak_ = *g*_leak_(*V* (*t*) − *E*_leak_). *I*_Na_ and *I*_K_ are:

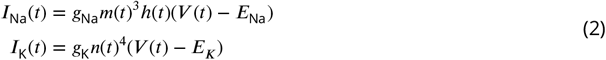

*I*_*base*_ is a constant input, simulating spontaneous ORN activity at rest without odorant stimulation (basal condition). *I*_noise_ = 1 nA ∗ *ξ* where *ξ* is Gaussian white noise with mean 0 and standard deviation 1. *I*_M_ is an M-type adaptation current given by:

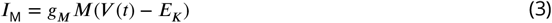

*I*_D_ simulates the exchange of ions between the dendrite and soma compartments and is given by *I*_D_ = *g*_VV_(*V* (*t*) − *V*_D_(*t*)). Each dendrite compartment, defined by its transmembrane potential *V*_D_, was modeled using:

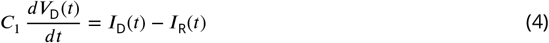

*I*_R_ is the receptor current, simulating the exchange of ions between the sensillar lymph and the dendrites through ligand-gated OR ion channels, given by:

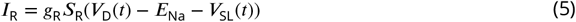

Dendrite conductivity, *S*_R_, depends implicitly on odor binding, activation and OR channel opening, approaching the steady state value *S*_R∞_ given by:

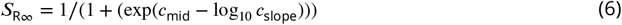

Odor binding, activation and channel opening are given in two stages by:

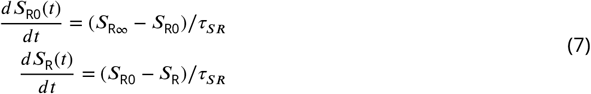

The potential of the sensillar lymph, defined by the local field potential *V*_SL_, is modeled by:

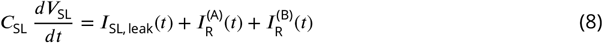

Here the sensillar leak current, simulating the flow of ions across the accessory cells, is given by

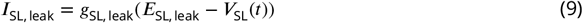

Each of the activation and inactivation variables *y*(*t*) = *n*(*t*), *m*(*t*), *h*(*t*), *M*(*t*) satisfy first-order kinetics:

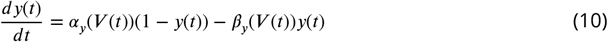

The equations for the nonlinear functions *α*_*y*_(*V*) and *β*_*y*_(*V*) were:

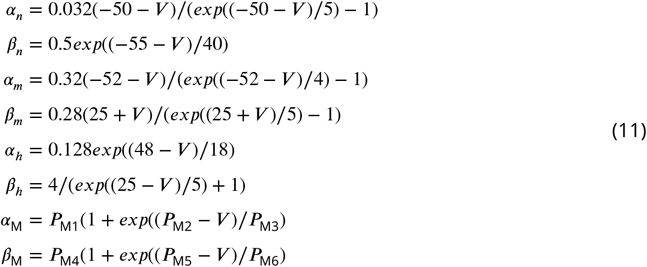

Receptor current (*g*_R_, *c*_mid_ and *c*_slope_) and M-current parameters (*g*_M_, *P*_M1_, *P*_M2_, *P*_M3_, *P*_M4_, *P*_M5_ and *P*_M6_) were fit to ab3A responses to methyl hexanoate dilutions using *scipy*.*optimize*.*minimize*.

Sensillum parameters (*g*_SL_ and *C*_SL_) were manually tuned to generate LFP and spike amplitudes comparable to published results (***Zhang et al., 2019; Nagel and Wilson, 2011; Martin and Alcorta, 2016***).

Parameters for B odorant sensitivity (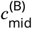 and 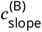) were determined manually by fitting to ab3B responses to 2-heptanone dilutions.

Morphometric differences between ORNs were taken into consideration by scaling B parameters for capacitance (*C* and *C*_1_) and conductance (*g*_leak_, *g*_Na_, *g*_K_, *g*_*VV*_, *g*_M_ and *g*_R_) by a scale factor of 0.74, factor representing the proportional surface area of B compared to A, calculated from the morphology parameters in ***Zhang et al. (2019***).

See **Table S2**) for specific parameter values.

Out computational model was implemented in Brian 2 version 2.6.0 (***Stimberg et al., 2019***) and model equations were integrated using the “milstein” method with Δ*t* = 0.05 ms timesteps.

The code is available at: https://github.com/tnowotny/drosophila-nsi/.

### Data analysis

Analyses and plotting were performed using custom scripts written in Matlab, unless otherwise specified. All plots and text report the mean of an experimental group ± standard error of the mean (SEM).

Pulse responses were defined as the average firing rate over a sampling window from 100 ms after valve opening to 100 ms after valve closing. These conditions are defined by the typical valve-to-response time, reflecting the stimulator-specific stimulus latency and stimulus-specific first spike latency.

Peri-stimulus time histograms were calculated as the average firing rate across trials in 10 ms bins smoothed by a 50 ms squared window. PID measurements in **Figure 9C** were averaged in the same way. Spike density plots were generated from custom scripts in Python, calculated as the average spike density function (Gaussian kernel with 10 ms window width) across trials.

Receptor and M current values in **Figures 5 and 7** were estimated by interpolating between spikes. This was done by determining the inflection point between each spike, defined as the near-zero minimum second derivative of the current trace. The second derivative was approximated using the Matlab *gradient* function.

### Experimental Design and Statistical Analysis

Spike response counts for female flies were defined as the number of A spikes within a 500 ms window starting from the release of the B odorant and were analyzed using Prism (GraphPad). The responses are reported and displayed as a firing rate. Where single comparisons between conditions were of interest, multiple t-tests were performed and adjusted using the Holm-Šídák correction **(Figures 1E and 2B)**. Where multiple comparisons between pulse timings were of interest **(Figures 2 and 3)**, a repeated measures ANOVA was performed with post-hoc analysis using Tukey’s test. Where comparisons between conditions and trends across measures were both of interest **(Figure 2)**, a Nested t-test was used to compare conditions with a Chi-square difference test across measures. Where comparisons between conditions were made with linear models **(Figures 2 and 3)**, Multiple linear regression was performed and tested with the sum of squares F-test.

**Figure 1.**
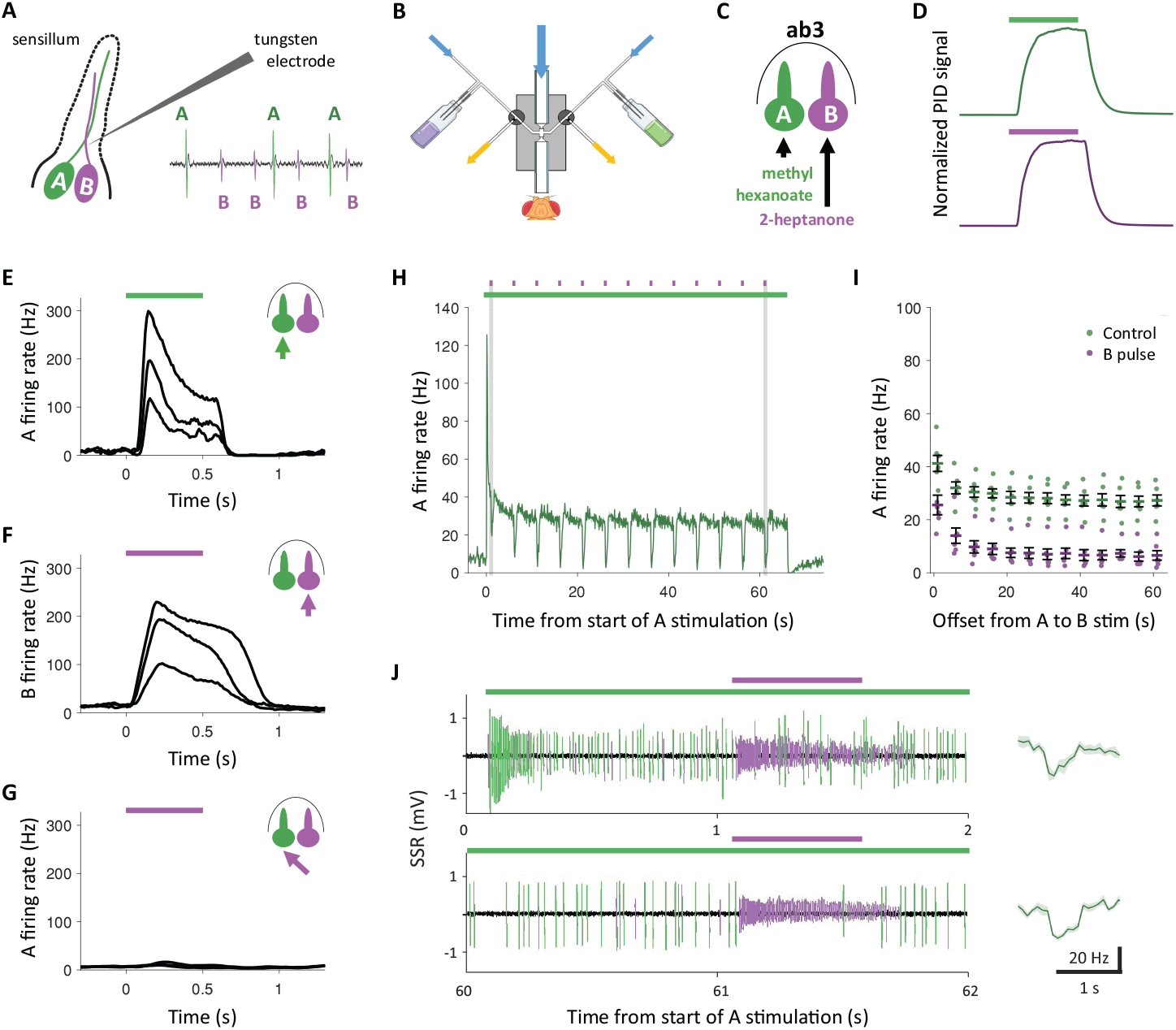
NSIs evident in ab3 during adaptation to sustained stimulation. **A)** An olfactory sensillum containing two ORNs (A and B) illustrating tungsten electrode placement during single sensillum recording (SSR) Inset: example SSR; ORN ‘A’ has a larger spike amplitude than ‘B’. **B)** Odor delivery apparatus: two odors delivered from separate valve-controlled airstreams join main air stream in a mixing block. Blue arrows = clean air entry. Yellow arrows = waste air to exhaust. **C)** The ab3 sensillum in which ORNs can be selectively stimulated by their indicated ‘cognate’ odors. **D)** PID signals normalized by peak response to 500 ms pulses of undiluted methyl hexanoate (green) and 2-heptanone (purple). **E)** Peri-stimulus time histogram (PSTH) of ab3A response to 500 ms pulse of cognate odor methyl hexanoate. **F)** PSTH of ab3B response to 500 ms pulse of cognate odor 2-heptanone. **G)** PSTH of ab3A response to 500 ms pulse of non-cognate odor 2-heptanone. Odors in E-G diluted in mineral oil at 10^−4^, 10^−5^ and 10^−6^. **H)** PSTH of mean ab3A response to background stimulation protocol: consecutive 500 ms B pulse stimulations (small purple bars above) along sustained 60 s A stimulation (long green bar). A stimulation 10^−5^ and B stimulation 10^−4^. Vertical shaded bars define time periods presented in panel J. **I)** Comparison of ab3A response to background stimulation protocol: A stimulation alone (control) and with B pulse. **J)** Sustained A stimulation only elicits a response from ab3A (large green spikes). A 500 ms B pulse stimulates ab3B (small purple spikes), which inhibits ab3A, 1 second after the start of A stimulation and 60 seconds later. Right: PSTH of mean response, shaded areas represent SEM.

**Figure 2.**
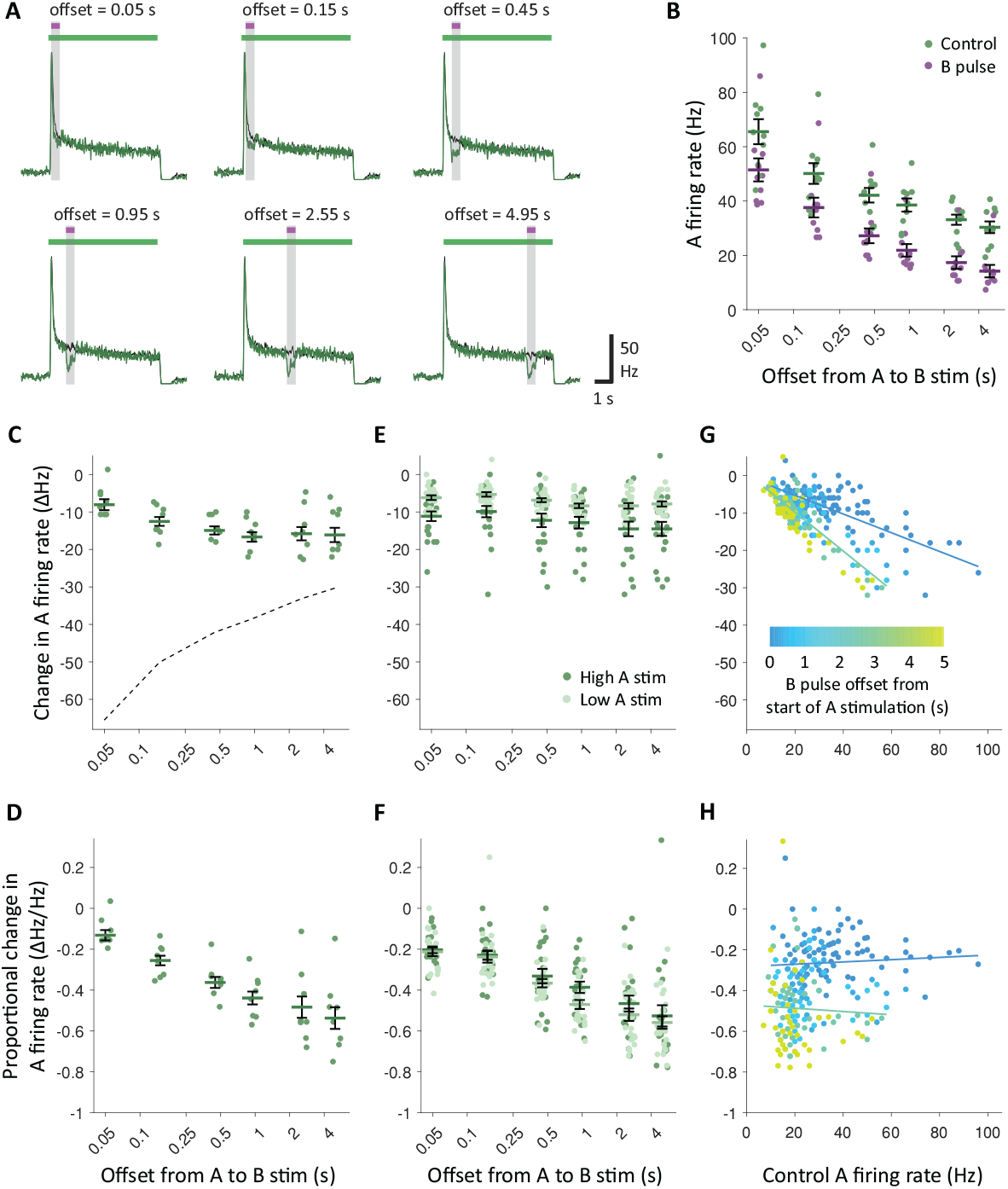
NSIs do not scale with adaptation-driven changes in ORN response. **A)** PSTH of mean ab3A response to fast stimulation protocol: 500 ms B pulses (purple bars) offset relative to A stimulations (green bars). Green trace shows A response during A and B stimulation; black trace shows response to A stimulation alone. A stimulation 10^−5^ and B stimulation 10^−4^. **B)** Comparison of ab3A response to fast stimulation protocol: A stimulation alone (control) and with B pulse. Here and in the following panels points represent individual ab3A responses and error bars represent group mean ± SEM. **C)** Relationship between offset of B pulse from start of A stimulation and change in ab3A firing rate from control (no B pulse) to trial (with B pulse). Dashed line delineates mean maximum possible inhibition, defined as the negative mean control firing rate. **D)** Relationship as panel C, with change in ab3A firing rate as a proportion of control firing rate. **E)** Relationship as panel C for high and low A stimulations during stimulation protocol. **F)** Relationship as panel E, with change in ab3A firing rate relative to control firing rate. **G)** Relationship between ab3A firing rate during control (no B pulse) and change in firing rate during trial (with B pulse). Colored according to offset of trial B pulse. Lines define linear regression of offsets during rapid adaptation (< 0.5 s - blue tones) and following (> 0.5 s - cyan to yellow tones). **H)** Relationship as panel G, with change in ab3A firing rate as a proportion of control firing rate. C and D, *n* = 10. E-F, Low A stim *n* = 21; High A stim *n* = 22.

**Figure 3.**
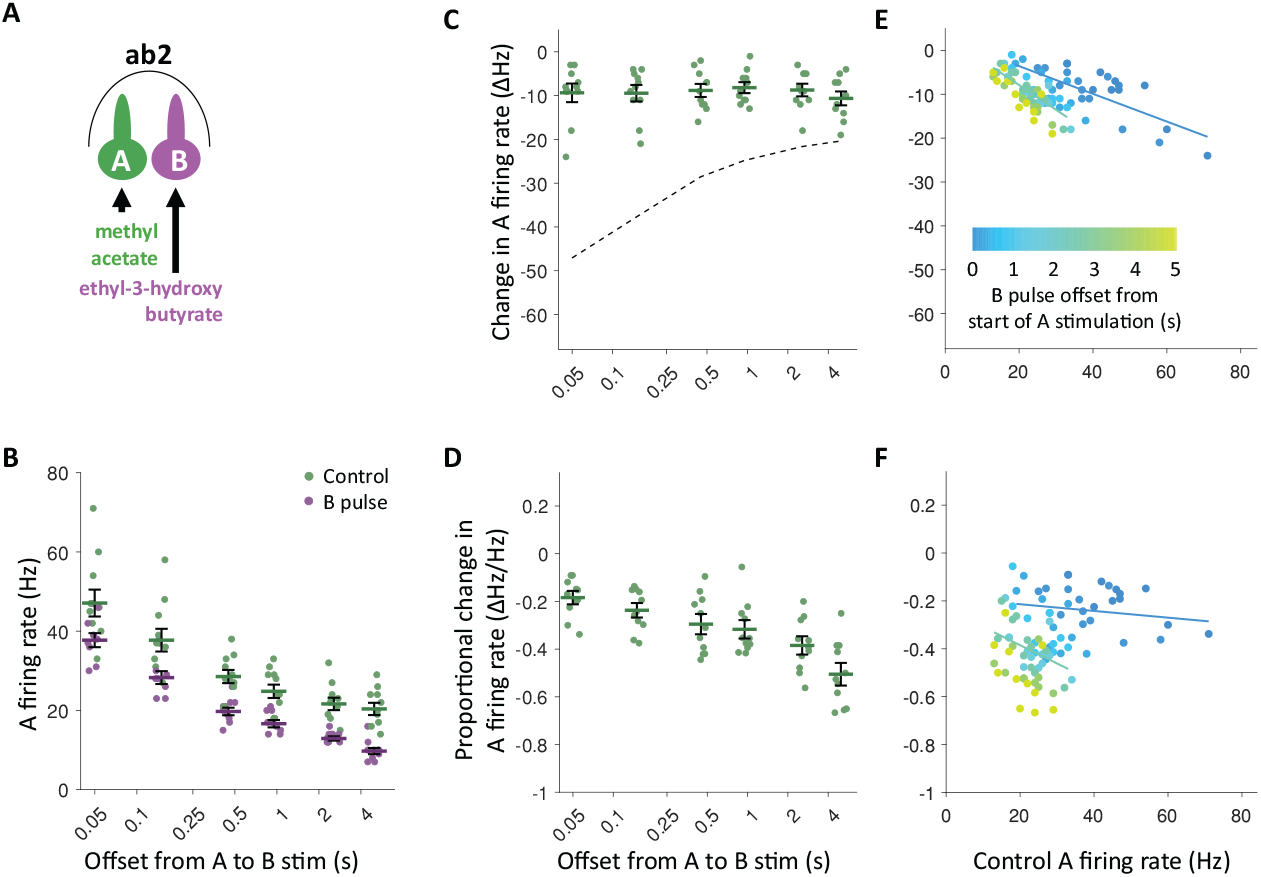
NSIs in ab2 also adaptation dependent. **A)** The ab2 sensillum in which ORNs can be selectively stimulated by their indicated ‘cognate’ odors. **B)** Comparison of ab2A response to fast stimulation protocol: A stimulation alone (control) and with B pulse. A stimulation 10^−4^ and B stimulation 10^−5^. Here and in the following panels points represent individual ab3A responses and error bars represent group mean ± SEM. **C)** Relationship between offset of B pulse from start of A stimulation and change in ab2A firing rate from control (no B pulse) to trial (with B pulse). Dashed line defines mean maximum possible inhibition (negative mean control firing rate) (*F*(3.260,32.60) = 1.889; *p* = .15; repeated measures ANOVA). **D)** Relationship as panel C, with change in ab2A firing rate as a proportion of control firing rate (*F*(3.503,35.03) = 28.23; *p* < .0001, repeated measures ANOVA; 0.05 s vs other offsets *p* < .03; Tukey’s multiple comparisons). **E)** Relationship between ab2A firing rate during control (no B pulse) and change in firing rate during trial (with B pulse). Colored according to offset of trial B pulse. Lines define linear regression of offsets during rapid adaptation (< 0.5 s - blue tones) and following (> 0.5 s - cyan to yellow tones). Data from panel C. (*F*(2,84) = 26.02; *p* < .0001; sum of squares F-test). **F)** Relationship as panel E, with change in ab2A firing rate as a proportion of control firing rate (*F*(2,84) = 23.57; *p* < .0001; sum of squares F-test).

Two-phase exponential models in **Figure S1** were generated with the Matlab *fit* function and their Euclidan distance compared with the *pdist* function.

Analytical comparisons of spike trains in **Figure 9** were completed in Python using the following functions: *numpy*.*linalg*.*norm* for the normalized Euclidian distance, *numpy*.*corrcoef* for the Pearson product-moment correlation coefficient and*elephant. spike_train_dissimilarity*.*victor_purpura_distance* for the Victor-Purpura distance.

Significance is reported as \**p* < .05, and data are expressed and graphed as mean±SEM. For one-way and two-way ANOVA, actual p values are reported when provided by the software. When actual p values are not provided by the software, only \**p* < .05, \*\**p* < .01, \*\**p* < .001, and \*\*\*\**p* < .0001 are reported.

## Results

### Non-synaptic interactions occur between co-housed ORNs during adaptation

ORN responses were measured via extracellular single sensillum recordings (SSRs) (**Figure 1A**). We examined B-to-A inhibition in the ab3 sensillum during variable offset dual-odor stimulations with a paradigm delivering two precisely timed odors, one selective for each ORN (**Figure 1B and 1C**). The odor that selectively activates the A ORN - its ‘cognate’ - was delivered alone or in combination with a short (500 ms) pulse of the odor that selectively activates the B ORN. The offset between the start of the A stimulation and the B pulse was varied to explore the relationship between stimulus timing and NSIs. By comparing the spiking activity of the A ORN with or without the B pulse, we determined the potential significance of NSIs under these conditions. Single odor photo-ionization detector (PID) measurements (**Figure 1D** show that both odors produce pulse shapes consistent with those reported previously (***Cafaro, 2016; Jordán et al., 2001; Martelli and Fiala, 2019***). As expected from their similar vapor pressures, the two odor pulses closely resemble each other and exhibit nearly identical arrival times. In addition, **Figures 1E and 1F** illustrate that ORNs respond robustly to their respective cognate odors at moderate dilutions, with **Figure 1G** showing no evidence of ab3A cross-activation by the cognate odor of ab3B.

We first examined NSIs during a long A stimulation (**Figure 1H**). We observed the characteristic pattern of adaptation: the A ORN initially responded with a rapid-onset high firing rate that then adapted – first quickly, then more gradually -— to a stable activity level ( 25% of the initial rate by 15-20 seconds). When pulses of the cognate odor for the B ORN were delivered on top of this A stimulation background, we observed high-magnitude NSIs at long delays, consistent with previous reports (***Su et al., 2012; Zhang et al., 2019***), with lateral B-to-A inhibition reducing the A firing rate to near the spontaneous activity levels observed before stimulation. For shorter offsets, we observed a relatively smaller but still significant reduction in the A firing rate, down to offsets as short as 1s (**Figure 1I and 1J**) (all *p* < .0006; *n* = 6; Multiple paired t-tests with Holm-Šídák correction. See **Table S1**) for test values). Therefore, if the A firing rate had not yet reached steady state before odor B was presented (offsets < 20 s), the activity during B stimulation – though reduced – remained above spontaneous activity.

### Non-synaptic interactions during short offset stimuli are of an effective lower magnitude

Having established that NSIs could be reliably reproduced, we next examined conditions more representative of natural odor plumes, where odor concentrations fluctuate on millisecond timescales rather than forming stable backgrounds (***Yee et al., 1995; Celani et al., 2014***). This extends previous detailed investigations of NSIs (***Su et al., 2012; Zhang et al., 2019***), which were conducted under stable “background odor” conditions, i.e. in the late phase of long A stimulations.

If NSIs played a significant role in the fast-fluctuation and short-onset-delay regime, we would expect their effects to be proportional to the firing rate of the inhibited ORN, which is considerably higher prior to adaptation than the stabilized firing rates during longer stimulations. In other words, we would expect the ‘effective’ inhibition (the change in firing rate as a fraction of the current firing rate) to remain comparable for both short and long onset delays. Prior work under ‘background odor’ conditions has already shown that the effects of NSIs strengthen if firing rates increase due to rising odor concentration (***Su et al., 2012***), meaning that in the ‘background odor’ regime and with respect to different odor concentrations, our expectation of roughly constant effective inhibition has already been demonstrated.

To directly assess NSI scaling for short onset delays, we delivered B pulses at offsets as short as 50 ms in spaced trials to eliminate residual effects from preceding stimuli (**Figure 2A**). We did not include shorter offsets or concurrent stimulations of both ORNs due to the insurmountable spike sorting challenges when both neurons have very high firing rates (see **Figure S2** and our earlier work on spike sorting (***Ellison et al., 2025***)). Across all tested offsets, we observed significant B-to-A inhibition (**Figure 2B** - all *p* < .00005; *n* = 10; Multiple paired t-tests with Holm-Šídák correction. See **Table S1**) for test values). However, contrary to our expectations, NSIs at short onset delays were not proportional to the A firing rate. Instead, NSI-mediated inhibition – quantified as the change in A firing rate during the B pulse – exhibited a slight inverse trend, increasing with larger B pulse offsets despite concurrent adaptation-driven reductions in the A firing rate (**Figure 2C**). Although repeated measures ANOVA demonstrated significant differences in absolute inhibition between offsets *F*(3.282, 22.97) = 7.765, *p* = .0007, *n* = 10), this increase was modest relative to the pronounced adaptation-induced decline in the A firing rate. Collectively, these findings suggest that NSI-mediated inhibition, as measured by absolute change in firing rate, remains broadly stable and largely independent of the ORN adaptation state. If anything, it is slightly increasing with A adaptation.

In order to analyze the *effective magnitude* of NSI-mediated inhibition for different ORN adaptation levels during short offset stimulations, we normalized the absolute reduction in firing rate due to NSIs by the (adaptation-dependent) control firing rate. We observed a roughly four-fold increase in the proportional reduction in A firing rate between 0.05 and 5-second offsets (**Figure 2D**). A Repeated measures ANOVA demonstrated differences between offsets (*F*(2.615, 18.31) = 24.11, *p* < .0001, *n* = 10). This relationship suggests that NSIs may be (relatively) less important for shorter odor onset delays, characteristic of natural mixed odor plumes. In order to assess how this observation interacts with the scaling of NSIs with stimulus intensity-driven firing rate differences in the inhibited ORN, we repeated short-offset stimulations across a range of A stimulation intensities. Due to limitations in precise concentration control in our olfactometer, trials were binned into ‘low’ and ‘high’ A stimulation intensity, defined by the initial peak response to the A stimulus when presented alone. Here we observed that higher A stimulation produced a greater reduction in the ab3A firing rate during B pulses (**Figure 2E**), consistent with stimulation-intensity-driven scaling of NSIs during ‘background condition’ (***Su et al., 2012***). A nested t-test revealed a significant difference in absolute NSI magnitude (change in A firing rate) between low and high A stimulation conditions (low: *n* = 21, high: *n* = 22, *t*(10) = 5.756, *p* = .0002). Notably, under these conditions, absolute NSI magnitude did not vary with B pulse offset at all (*p* = .28; Chi-square test; X^2^(1) = 1.17), supporting the conclusion that NSIs lead to broadly the same absolute change in firing rate independent of stimulus timing and hence the adaptation state of the inhibited ORN. When data were normalized to assess effective inhibition (proportional firing rate change) we again observe an increase at larger B pulse offsets **(Figure 2F)** (*p* < .0001; Chi-square test; X^2^(1) = 117.5). However, no significant difference in effective inhibition was found between low and high A stimulation groups (nested t-test, *t*(10) = 0.3806, *p* = .71). These observations indicate that the dependence of NSIs on the inhibited neuron’s firing rate is not the same if this firing rate is changed by stimulus concentration or by adaptation.

To illustrate this discrepancy more directly, we plotted the change in A firing rate directly relative to the A control firing rate, regardless of the B pulse offset. The pulse offset, and hence A adaptation state, was then indicated by color-coding the data points (**Figures 2G and 2H**). When examining the absolute change in firing rate, (**Figure 2G**), we clearly see that data points form two clusters, one for offsets < 0.5 s (blue) corresponding to the period of most rapid A adaptation (see Figure 1F) and another for offsets > 0.5 s (cyan to yellow) when A is largely adapted. To examine the explanatory power of these two clusters, the data were fit to separate linear regression models, which were found to fit the data better than a single model (*F*(2,254) = 108.8, *p* < .0001; sum of squares F-test). The resulting linear models both intercept 0 ΔHz near 5 Hz, but differ in slope, with adapted (> 0.5 s) offsets leading to larger changes in the A firing rate as the A stimulation intensity increases (slope −0.47 vs −0.24 for < 0.5 s offsets). When examining effective inhibition (**Figure 2H**) we noted the same pattern of two data clusters corresponding to the periods of rapid adaptation and following slow adaptation/ steady state. We again analyzed these two clusters using separate linear models, which again fit the data better than a single model (*F*(2,254) = 58.47, *p* < .0001; sum of squares F-test). However, in this presentation, the resulting linear models have near-zero slope, consistent with the pattern that effective NSI-mediated inhibition is constant across A stimulation intensity. The models for the two clusters instead differ in intercept, with adapting (> 0.5 s) offsets leading to larger proportional changes in the A firing rate (intercept −0.45, vs −0.21 for < 0.5 s offsets; slope −0.0003 vs −0.0002).

Overall, these results indicate that – unlike differences in A firing rate due to stimulation intensity, where NSI magnitude scales reliably with ORN firing rate – differences in A firing rate due to varying adaptation state do not unambiguously predict NSI magnitude. Instead, even when starting from the same A firing rate, one can observe different levels of NSI-mediated inhibition, both in absolute and relative terms. This demonstrates clearly that the impact of NSIs on the inhibited ORN’s firing rate depends on its adaptation state, with strong effective inhibition occurring only at large onset offsets, similar to those used in most previous studies. In contrast, during the short-offset stimuli examined here (when B stimulation follows A stimulation by only tens of milliseconds), minimal adaptation and high initial firing mean that the same absolute change in firing rate produces proportionally weaker NSIs.

We have confirmed these results in a second sensillum type (**Figure 3**) and further demonstrated that the pattern is recapitulated as A recovers from adaptation following a prolonged high-intensity stimulation (**Figure S1**).

### A spiking electrical circuit model replicates the adaptation dependence of effective NSIs

Given that NSIs are electrical in nature and thus occur on rapid timescales, their observed dependence on adaptation appeared difficult to explain through reasoning alone. Why would an essentially instantaneous interaction need long adaptation times to become relevant? Further experimental investigation of this finding is constrained by the limitations of SSRs and the practical inaccessibility of the ORN soma for intracellular recordings. To address this, we turned to computational modeling to explore whether and how the electrical architecture of the sensillum could give rise to adaptation-dependent NSIs.

The understanding of NSIs has been significantly influenced by computational modeling (***Vermeulen and Rospars, 2004; Su et al., 2012; Zhang et al., 2019***). However, existing models offer a limited exploration of the electrical interactions between ORNs, because they focus on asymptotic or steady-state solutions, which do not account for the full temporal evolution of responses or the dynamics of adaptation, and may therefore overlook important aspects of the underlying electrical interactions. To address these limitations, we developed a spiking model of two co-housed, two-compartment, Type 1 Hodgkin-Huxley neurons with adaptation modeled by an M-type spike-frequency adaptation current. The neurons share common dendritic and somatic extracellular spaces, which have separate extracellular potentials. When the neurons are activated by odors and change their conductance, the voltage difference between the extracellular spaces changes, which in turn changes the reversal potential of the receptor currents of both neurons, and so causes inhibition through a reduced “driving force” (see the role of the sensillum potential (local field potential) *V*_SL_ in equation (5)). The model, conceptually summarized in **Figure 4A**, produces ORN responses with realistic electrical changes (**Figure 4B**. In order to enable a naive examination of NSIs, the model was only minimally tuned to single ab3A odor responses (**Figure S3**). Additional features, such as LFP amplitudes and ab3B size, have been scaled to available data in order to generate appropriate output (see Methods). We first examined how well our sensillum model reproduces the general features of NSIs. We began by comparing overall response characteristics by analyzing the time-resolved ORN activity during B-pulse stimulations overlaid on A background stimulations, paralleling the SSR experiments in Figure 2. As simulations report individual ORN activity and therefore negate the requirement for spike sorting, we were able to include fully concurrent stimulation of both ORNs – an aspect not accessible in our experimental recordings. Simulated responses closely mirror SSR data, with clear lateral inhibition of A background activity during B pulses **(Figure 4B and 4C)**. Relative to the SSR recordings, simulated NSIs are more pronounced and more dynamic, particularly for B pulses delivered later in the A stimulation when A is strongly adapted. At the onset of B stimulation, the shared LFP exhibits a sharp decrease, thereby reducing the effective driving force on A, which in turn lowers the receptor current and leads to a drop in firing rate. Although the LFP remains relatively stable for the remainder of the B pulse, the A firing rate gradually recovers and, upon termination of B stimulation, a marked rebound in A firing is observed. Both features indicate substantial de-adaptation in response to the sustained reduction in receptor current induced by B stimulation.

**Figure 4.**
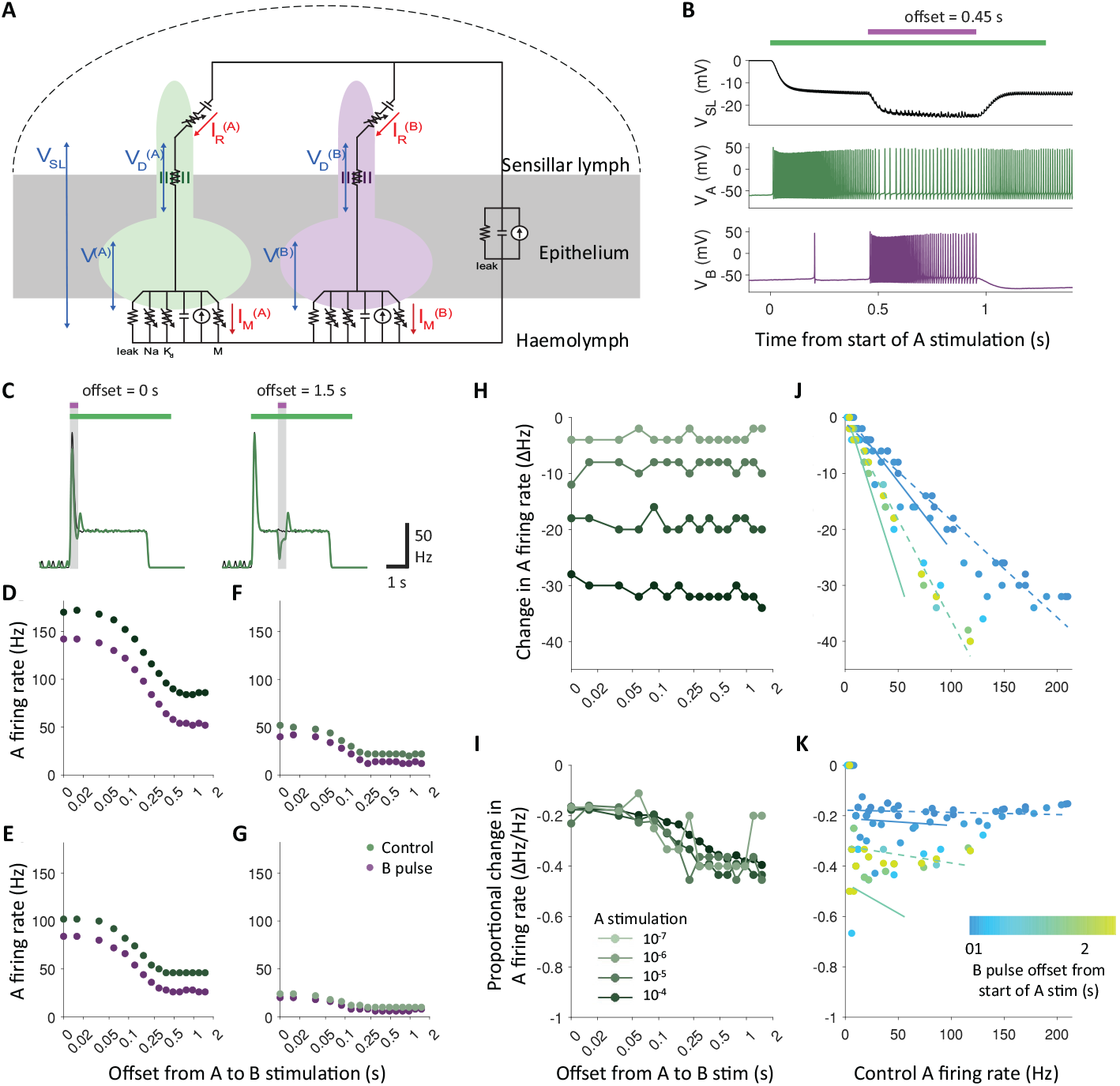
A spiking electrical circuit model of compartmentalized insect ORNs replicates adaptation-dependence of NSIs. **A)** Simplified diagram of electrical circuit model of an olfactory sensillum consisting of two ORNs (A and B) and an auxiliary cell (gray rectangle). Relevant currents (red) and potential differences between compartments (blue). **B)** LFP (V_SL_) and soma potentials for A (green) and B (purple) during simulated experiment. **C)** Spike density function of A response to simulated experiment: 500 ms B pulses (purple bars) offset relative to A stimulations (green bars). Green trace shows A response during A and B stimulation; black trace shows response to A stimulation alone. Vertical shaded bars define sampling windows. Comparison of simulated A response during stimulation protocol: A stimulation alone (control) and with B pulse. **D)** 10^−4^, **E)** 10^−5^, **F)** 10^−6^ and **G)** 10^−7^. B stimulation 10^−4^ in all panels. **H)** Relationship between offset of B pulse from start of A stimulation and change in simulated A firing rate from control (no B pulse) to trial (with B pulse). **I)** Relationship as panel H, with change in A firing rate as a proportion of control firing rate. **J)** Relationship between A firing rate during control (no B pulse) and change in firing rate during trial (with B pulse), colored according to offset of trial B pulse. Dashed lines define linear regressions for the data presented; Solid lines for the SSR data from Figure 2. Two linear regressions calculated for offsets during rapid adaptation (< 0.5 s - blue tones) and following (> 0.5 s cyan to yellow tones). **K)** Relationship as panel J, with change in A firing rate as a proportion of control firing rate.

Our second objective was to evaluate whether the electrical model sensillum replicates the adaptation-dependent modulation of NSIs observed during short offset odor stimuli. Due to the implementation of a single adaptation element in the model (yielding response stabilization within approximately 1 s), we restricted our analysis to B pulse offsets less than 2 s, reasoning that longer delays would contribute minimal additional information. In addition to the inclusion of fully concurrent stimulation of both ORNs, we also used a more granular stimulation protocol, incorporating a denser sampling of B pulse offsets over this reduced range to more precisely characterize NSI evolution relative to the adaptation state of the inhibited ORN.

Overall, the pattern of simulated NSIs closely resembled the SSR experiments **Figure 4D-4I**. B pulse stimulation consistently suppressed A firing across all tested offsets, with the magnitude of inhibition increasing at higher A stimulation intensities. Inhibition magnitudes are also quantitatively comparable to those observed experimentally, with similar degrees of absolute inhibition (change in a firing rate), which remains largely independent of B pulse timing. When normalized to reflect effective inhibition (proportional change in A firing rate), simulated responses continue to align with SSR data. Under this metric, differences driven by A stimulation intensity are eliminated, and a uniform increase in effective inhibition is observed with increasing B pulse offset. These findings support the conclusion that while absolute inhibition is governed by stimulation intensity of the inhibited ORN, effective NSI strength is primarily determined by adaptation state. As with SSR data, simulated results were also analyzed versus A control response **(Figure 4J and 4K)** in order to compare the relationship between stimulation intensity, adaptation state and NSI magnitude. We again observed similar results, with the formation of two distinct data clouds corresponding to early (< 0.5 s) and late (> 0.5 s) adaptation of A, with longer B stimulation offsets leading to higher magnitude NSIs.

Our model, being purely electrical and tuned only to single ORN responses, closely replicates experimental findings, supporting an electrical basis for the adaptation dependence of NSIs. Since the model is driven by just two dynamic inputs – receptor (R) current and M-type adaptation current – NSI effects should be predicted by a composite ‘net input current’ representing their combined influence. To test this, we analyzed R and M currents across B pulse offsets to compute net input current over time.

As expected from a conductance-based model, simulated currents include changes due to spiking activity **(Figure 5A-5C)**. As modeled R and M currents originate in separate compartments (dendrite and soma, respectively), visible spikes are of a differing magnitude. Therefore, in order to allow for a comparison of the relative effect of the currents, we estimated the effective currents besides the effects of spiking (see Methods for details). We compared these estimated currents as well as the net input current (absolute value of R - M) during A stimulation alone and with B stimulation **(Figure 5D and 5E)**. The R current changes more rapidly, consistent with receptor activation kinetics, while M current increases are more gradual, reflecting the slower kinetics of spike-frequency adaptation. During B pulses, both currents are reduced by NSIs. These reductions are interpreted as follows: R current decreases result from LFP-mediated attenuation of the driving force, while M current decreases reflect de-adaptation following from a reduction in spiking.

**Figure 5.**
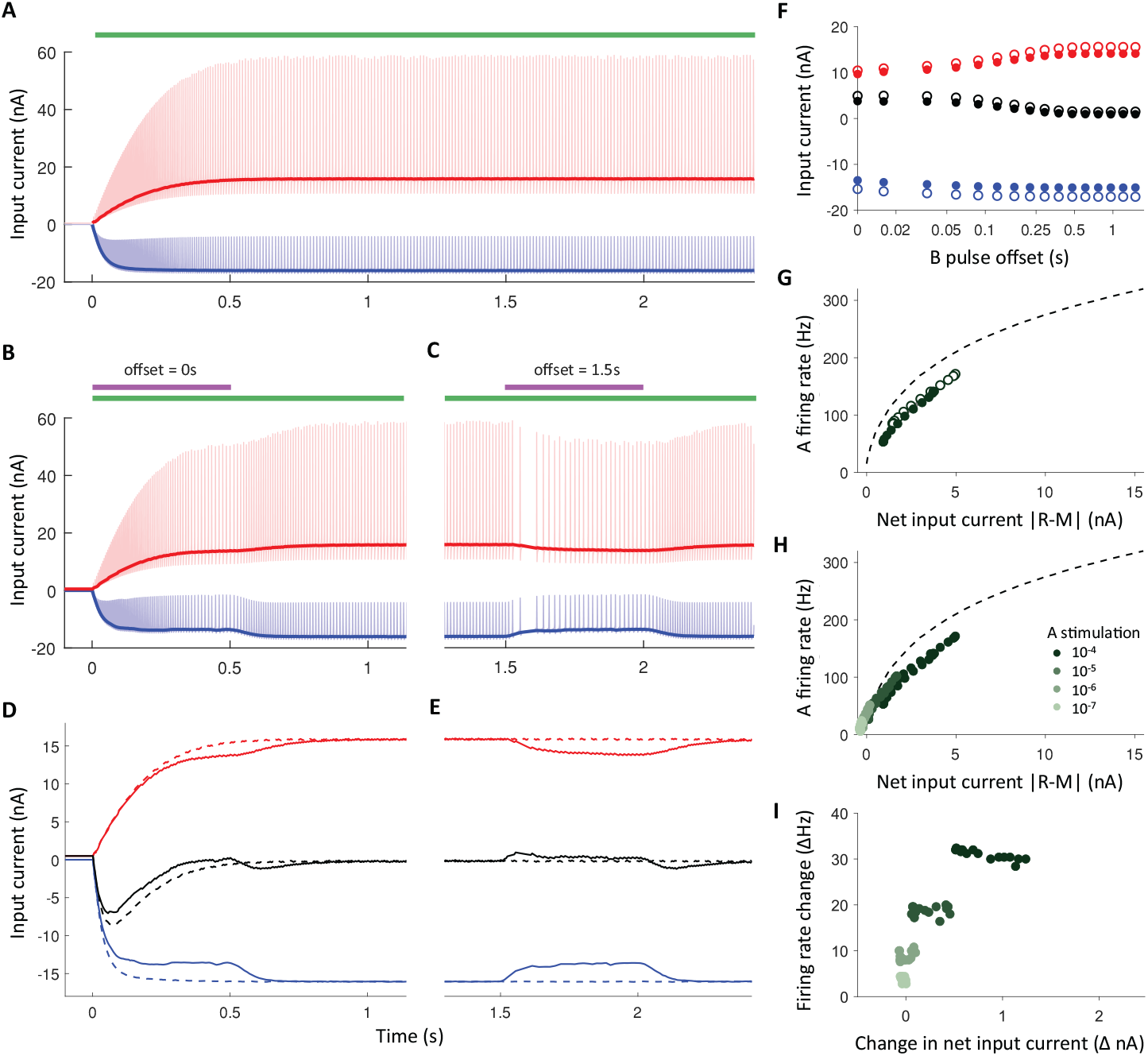
Simulated electrical interaction of adaptation and NSIs. Time-resolved simulation of A receptor (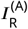 blue) and M-type adaptation currents (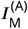 red) during A stimulation alone (green bar). Lines illustrate estimated effective current values behind spiking. **B)** As panel A during experimental trial with concurrent stimulation of A and B (purple bar). **C)** As panel B following long offset before B stimulation. **D)** Estimated currents during concurrent stimulation of A and B, including Black = net input current |*R* – *M*|. Dashed lines = A stimulation alone; solid lines = with B pulse. **E)** As panel D during long offset of B stimulation. **F)** Simulated ORN currents during A stimulation alone (hollow) and with B pulse stimulation (filled) at increasing B stimulation offsets. **G)** Relationship between net input current and A firing rate during A stimulation alone (hollow) and with B pulse stimulation (filled). Dashed line = frequency-current (F-I) curve generated from current injection steps. **H)** Relationship between net input current and A firing rate at a range of A stimulation intensities. Both A stimulation alone and with B pulse represented by filled points. **I**) Relationship between change in net input current from control to trial and change in A firing rate at a range of A stimulation intensities. All B stimulations 10^−4^. A stimulation 10^−4^ in panels A-G; B stimulation 10^−4^ in all panels

We next compared the average effective current values across each 500 ms sampling window for the range of B pulse offsets **(Figure 5F)**. During A stimulation alone, the R current increases slightly at later time points, reflecting full receptor activation. The M current also increases, although more slowly and significantly, following the A adaptation time course. This leads to a comparable reduction in the combined net input current over time during A stimulation alone. When B pulses are also delivered, NSIs lead to a consistent reduction in the R current, independent of B pulse offset. This is expected as B stimulation intensity is unchanged, leading to the same LFP deflection. In contrast, the M current is not equitably impacted by NSIs, with a greater reduction occurring during longer B pulse offsets. These changes combine, leading to reduced differences in the net input current due to NSIs at longer B pulse offsets as the A adaptation state progresses.

In order to examine the significance of net input current at increasing B pulse offsets, we compared these values (and the corresponding A firing rates) to the model frequency–current (F–I) curve, derived from simulations of constant direct current injections **(Figure 5G and 5H)**. The values are consistent with the F-I curve, supporting the robustness of our effective current estimation method. Furthermore, when we examine the change in net input current induced by NSIs during B pulses alongside the corresponding change in the A firing rate **(Figure 5I)** we observe that, although net input current is reduced less by longer B pulse offsets, the resulting suppression of A firing remains relatively constant (as described previously in both SSR and simulated data). We understand this pattern to be consistent with the non-linear shape of the F-I curve. During high stimulation intensity and early adaptation, net input current is high and the slope of the F-I curve is shallower; therefore, current changes lead to smaller firing rate changes. As the M current increases during late adaptation, net input current falls, the slope of the F-I curve increases, and the same change in current leads to larger firing rate changes.

Given that both the overall pattern of NSIs and their temporal dynamics appear to depend on the adaptation state of the inhibited ORN, we suggest that dynamic adaptation (including deadaptation) interacts with NSIs in a non-linear and therefore somewhat non-intuitive manner. We propose that, in the absence of dynamic adaptation, NSIs would lead to a constant current change, following from the stimulus-timing independent change in the driving force. The effect of NSIs would then essentially only depend on the F-I curve of the receptor neuron, with stronger effects at low firing rates (steep F-I curve) and less strong effects at high firing rates (flatter F-I curve).

### Adaptation dependence of NSIs explained by non-linear electrical interactions

To investigate the non-linear effects of dynamic adaptation during NSIs, and to assess whether our expectations for a more direct dependence on the F-I curve could be recovered, we conducted additional simulations. In these simulations, the adaptation state of the A ORN was kept constant during each B pulse by not simulating the dynamics of the M current, thereby preventing any further adaptation, or de-adaptation, from taking place **(Figure 6A)**. The remaining model parameters were the same as those used previously, simulating the odor response and morphology of the ab3 ORNs.

**Figure 6.**
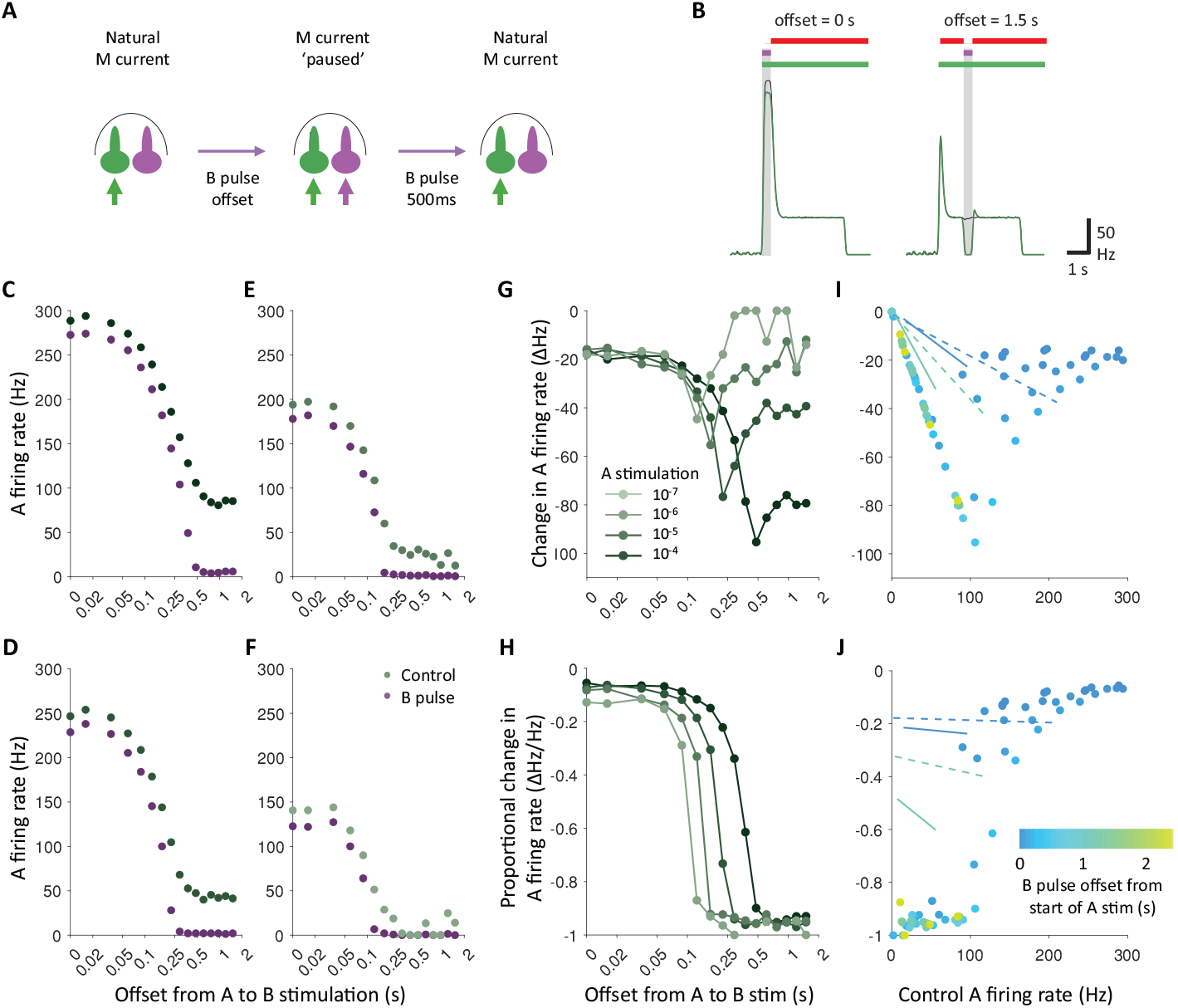
Controlling adaptation increases adaptation-dependence of NSIs. **A)** Experimental design. **B)** Spike density function of A response to simulated experiment with adaptation dynamics paused during B stimulations: 500 ms B pulses (purple bars) offset relative to A stimulations (green bars). Gaps in red bars indicate periods when the M-type adaptation current was paused. Green trace shows A response during A and B stimulation; black trace shows response to A stimulation alone. Vertical shaded regions indicate sampling windows. Comparison of simulated A response during stimulation protocol: A stimulation alone (control) and with B pulse. **C)** 10^−4^, **D)** 10^−5^, **E)** 10^−6^ and **F)** 10^−7^. B stimulation 10^−4^ in all panels. **G)** Relationship between offset of B pulse from start of A stimulation and change in simulated A firing rate from control (no B pulse) to trial (with B pulse). **H)** Relationship as panel G, with change in A firing rate as a proportion of control firing rate. **I)** Relationship between A firing rate during control (no B pulse) and change in firing rate during trial (with B pulse), colored according to offset of trial B pulse. Solid lines define linear regressions for the SSR data from Figure 2; Dashed lines for simulated data from Figure 4. **J)** Relationship as panel I, with change in A firing rate as a proportion of control firing rate.

**Figure 7.**
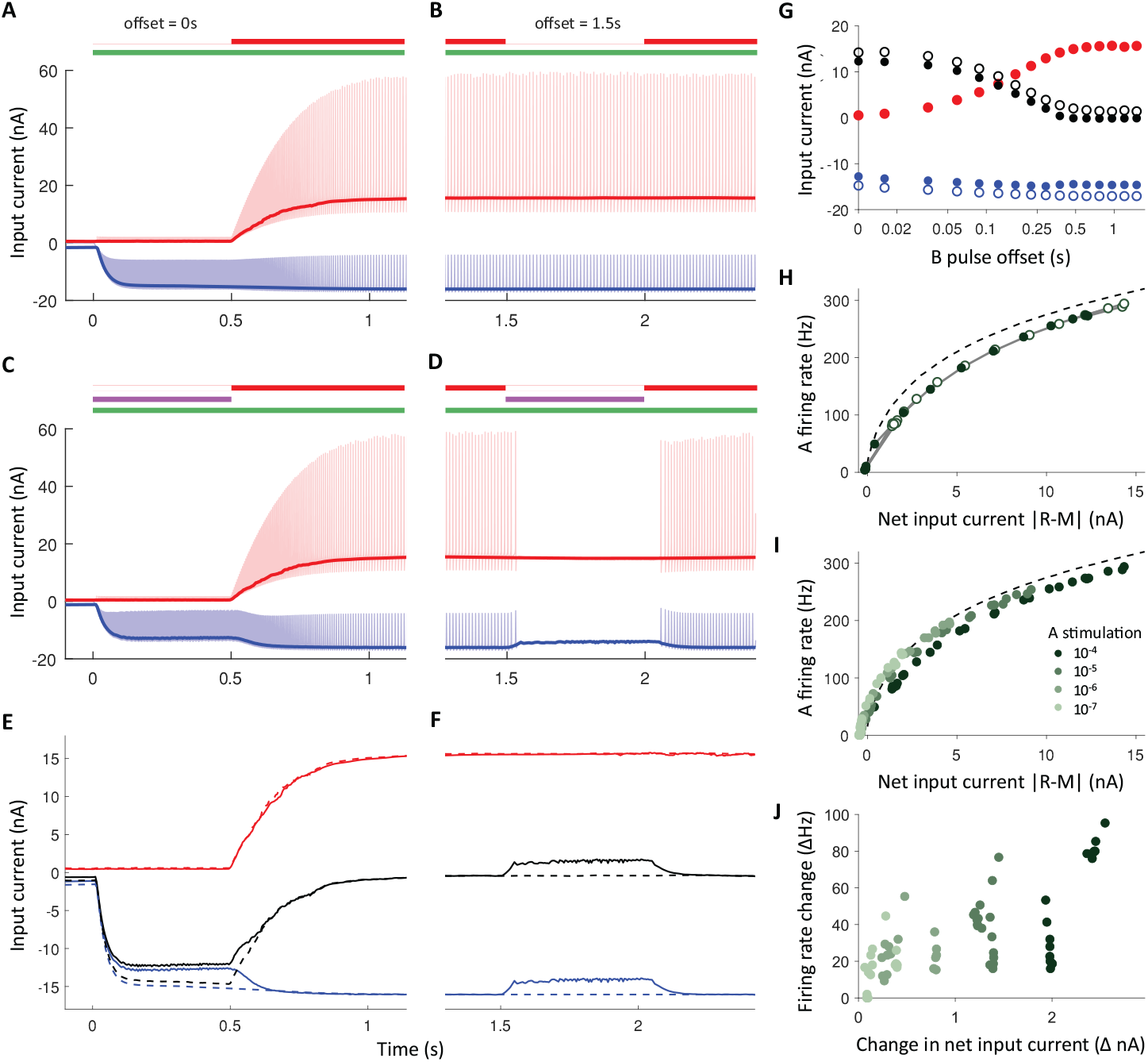
Simulated electrical interaction of adaptation and NSIs during controlled adaptation. **A)** Time-resolved simulation of A receptor (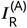 - blue) and M-type adaptation currents (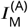 - red) during A stimulation alone (green bar), with M current dynamics paused during the first 500 ms of stimulation (gap in red bar), serving as a control condition. Lines illustrate estimated effective current values behind spiking. **B)** As panel A, with M current paused for 500 ms following a long offset. **C)** As panel A during experimental trial with concurrent stimulation of A and B (purple bar) during paused M current dynamics. **D)** As panel B following long offset before B stimulation and M current pause. **E)** Estimated currents during concurrent stimulation of A and B, including Black = net input current |*R* – *M*|. Dashed lines = A stimulation alone; solid lines = with B pulse. **F)** As panel E during long offset of B stimulation. **G)** Simulated ORN currents during A stimulation alone (hollow) and with B pulse stimulation (filled) at increasing B stimulation offsets. **H)** Relationship between net input current and A firing rate during A stimulation alone (hollow) and with B pulse stimulation (filled). Dashed line = frequency-current (F-I) curve generated from current injection steps. **I)** Relationship between net input current and A firing rate at a range of A stimulation intensities. Both A stimulation alone and with B pulse represented by filled points. **J)** Relationship between change in net input current from control to trial and change in A firing rate at a range of A stimulation intensities. All B stimulations 10^−4^. A stimulation 10^−4^ in panels A-G; B stimulation 10^−4^ in all panels.

In the “constant M current” condition, A firing rates during sampling windows are more stable: adaptation does not proceed at short B pulse offsets and de-adaptation is absent at longer offsets **(Figure 6B)**. Lateral inhibition of A activity remains evident across all B pulse offsets at a range of A stimulation intensities **(Figure 6C-6F)**.

However, in contrast with natural ‘dynamic’ adaptation conditions, NSI magnitude does not consistently scale with A stimulation intensity **(Figure 6G-6J**). At longer B pulse offsets, the originally observed scaling is preserved, NSIs are very strong and A activity is effectively halted during B stimulation. However, at shorter B pulse offsets, absolute inhibition (change in A firing rate) does not scale with A stimulation intensity, leading to a slight increase in effective inhibition a lower stimulation intensities. Overall, NSIs display a greater adaptation-dependence without dynamic adaptation.

Our findings indicate that dynamic adaptation during NSIs in fact reduces absolute NSI-mediated inhibition at longer offsets, counteracting the basic effect of adaptation through reduced further adaptation (small offsets) or de-adaptation (large offsets). Therefore, in the absence of dynamic adaptation, effective inhibition increases even more significantly with adaptation state.

In the final analysis of adaptation-controlled simulations, we again analyzed R, M and net input currents across B pulse offsets. In comparison to A stimulation alone **(Figure 7A and 7B)**, in the stabilized M-current regime the addition of B stimulation reduced only the R current **(Figure 7C and 7D)**. This selective reduction, unposed by the dynamic M current, leads to a larger change in the net input current **(Figure 7E and 7F)** compared to results in Figure 5 with natural M current dynamics.

When comparing the average effective current values across each 500 ms sampling window for the range of B pulse offsets, net input current reduces more substantially as the A adaptation state progresses **(Figure 7G)**. However, the reduction in net input current due to lateral inhibition, being exclusively driven by changes in the R current, is constant and independent of B pulse offset. When compared with simulations incorporating dynamic adaptation (Figure 5), the modulation pattern of the R current remains broadly consistent. Notably, there is a slight increase in R current suppression during NSI conditions (trial vs control), likely attributable to the fixed M current, which does not decline during trials. This maintains a more hyperpolarized membrane potential and consequently increases the driving force and subsequent R current. In contrast, the M current displays more substantial differences: while it remains comparable at longer offsets, it is significantly reduced at shorter offsets. These combined effects produce a markedly higher net input current at short offsets, consistent with the elevated firing rates observed in these conditions.

As in dynamic adaptation simulations, a comparison of net input current against A firing rate reveals a relationship consistent with the model F–I curve **(Figure 7H and 7I)**. However, here a wider range of currents and firing rates are described. This pattern is consistent with the moderating role of adaptation, generally serving to adjust sensory responses to maintain a narrow range of firing rates and prevent saturation due to strong stimulation.

When examining the changes in net input current induced by NSIs alongside the corresponding changes in the A firing rate **(Figure 7J)**, we observe a more constant current change at a fixed A stimulation intensity. This finding is consistent with offset-invariant membrane conductivity, which is determined by the proportion of open receptor channels resulting from the constant intensity of the stimulation. During periods of rapid adaptation (B pulse offsets < 0.5 s), the net input current remains most stable. At the highest A stimulation intensity (10^−4^), these early offsets consistently show current changes of approximately 2 nA and associated firing rate changes below 60 Hz. In contrast, longer offset durations are associated with larger and more variable reductions in net input current and higher firing rate changes, reflecting reductions of both to zero. At lower stimulation intensities, this pattern becomes less distinct, likely due to an increased influence of model noise under these conditions, as noted previously.

Overall, we find that by eliminating dynamic adaptation during NSIs we recover a relatively constant change in net input current. This constant change leads to absolute firing rate changes that increase substantially with increasing adaptation state. This contrasts with dynamic adaptation, where absolute firing rate changes due to NSIs are constant and the change in net current varies. However, both of these patterns are consistent with the model F-I relationship. Due to the non-linearity of this relationship, the same current change leads to small changes in firing rate during early adaptation (when net input current is high) and larger changes during late adaptation (when net input current is comparatively low).

Based on these findings, we conclude that the relatively constant absolute inhibition observed under conditions of dynamic adaptation arises from the inherently normalizing influence of adaptation mechanisms. During early adaptation, slightly higher levels of absolute NSI-mediated inhibition are observed. This is attributable to reductions in net input current, driven by an increasing M current, which shift neuronal responses into a steeper region of the F–I curve, thereby amplifying the impact of current changes on firing rate. In contrast, during later adaptation phases (when the M current has stabilized at a high level), absolute NSIs are substantially reduced due to compensatory adaptation dynamics. NSI-induced reductions in R current produce corresponding decreases in firing rate, which then trigger de-adaptive reductions in the M current and a subsequent partial recovery in firing rate. Consequently, the modulatory effects of adaptation act to suppress NSI-mediated reductions in firing during this later phase.

Overall, our experimental and modeling results demonstrate that higher magnitude effective NSIs only occur when A is in a later adaptation state. Like experimental data, simulated data form a distinctive pattern where B stimulation of the same intensity leads to roughly the same reduction in the A firing rate, independent of B stimulation timing. When this change is considered along with the rapid reduction in the A firing rate taking place due to adaptation, only longer offsets of odors lead to a significant effective inhibition (proportional change in the A firing rate). From this, we conclude that NSIs are less significant during short offset odor stimuli, making them an unlikely mechanism for interpreting temporal features in odor mixtures, contrary to what was previously suggested (***Pannunzi and Nowotny, 2021; Bandyopadhyay and Sachse, 2023***).

Mechanistically, we attribute this reduction to the previously unexamined influence of adaptation currents operating in ORNs, summarized in **Figure 8**. Although activation of one co-housed ORN can reduce the driving force on another, this second ORN will only display NSIs if certain conditions are met. Firstly, strong NSIs only occur when ORNs are strongly stimulated. NSIs result from changes to the driving force via changes to the LFP. ORNs are maximally sensitive to these changes during strong stimulation when a large proportion of receptors are open and conductivity is high. Secondly, strong NSIs can only occur when ORNs are strongly adapted. Without adaptation currents to oppose the high receptor current, ORNs are close to their maximum firing rate and hence less sensitive to input current changes (described by the F-I curve). When stimulation and adaptation are strong, receptor and adaptation currents combine to form a lower input current, moving to response conditions where even a small change in current results in a large change in activity. Finally, the effect of adaptation continuously interacting with firing rate in a closed-loop means that reduced adaptation or even de-adaptation during NSIs reduces the visible effects of NSIs somewhat.

**Figure 8.**
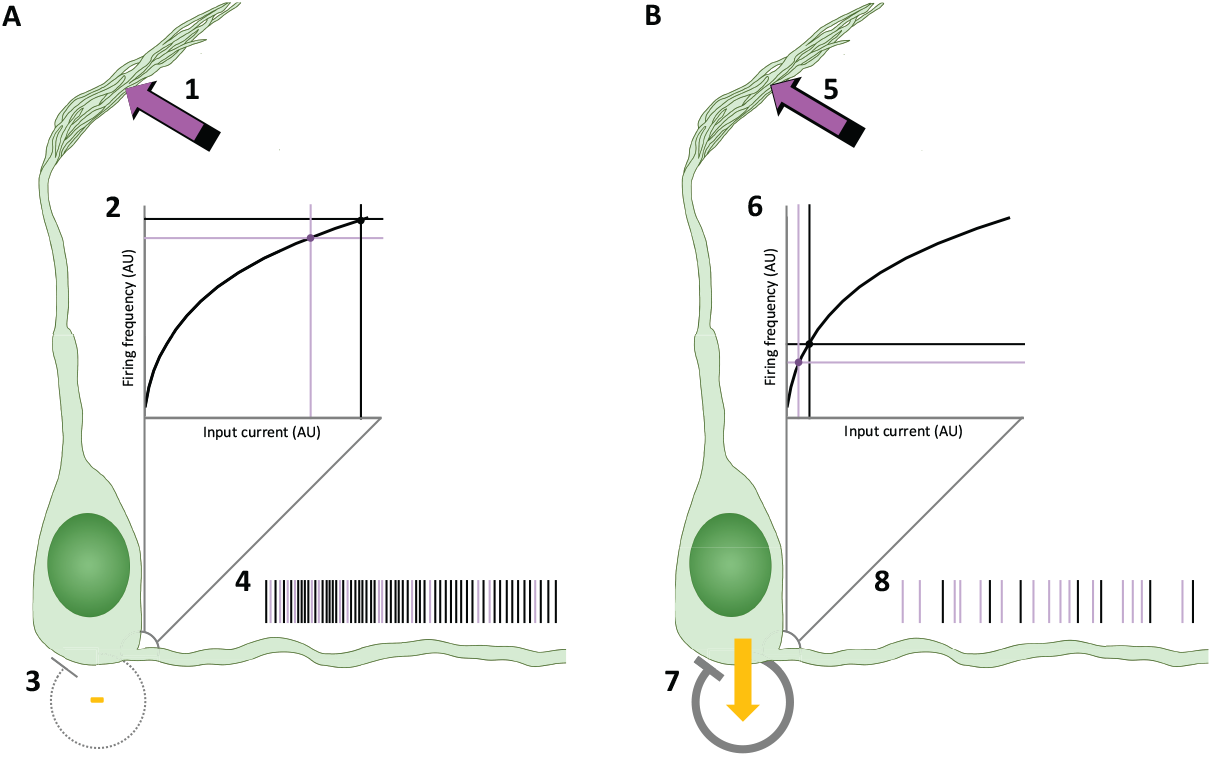
NSIs during early and late adaptation. When odors fluctuate rapidly **(A)** and A and B are stimulated together, NSIs reduce the driving force (1 – purple arrow) in comparison to A stimulation alone (black arrow). The resulting receptor current forms the input current driving the firing response at the spike initiation zone (2). As the receptor current is not modified by an adaptation current (3), the overall input current is high and, according to the shallow F-I relationship at high stimulations, NSIs lead to a proportionally small drop in firing (4). When odors fluctuate more slowly **(B)**, NSIs reduce the driving force to the same extent (5), however, the firing response (6) is now modified by a strong inhibitory adaptation current (7). Receptor and adaptation currents combine to an overall lower input current and hence firing activity governed by a steeper F-I relationship. Therefore, current changes due to NSIs lead to a proportionally large drop in firing (8). Purple spikes represent those lost during NSIs.

In summary, NSIs are high when the inhibited neuron is adapted because the low firing rate (steep F-I curve) coincides with high receptor conductance, allowing for large changes of input current when the LFP changes due to B activity. This persists against partial masking by de-adaptation. In other situations, where either the receptor conductance is low, or where the receptor conductance is high but the firing rate is also high due to minimal adaptation, NSIs are comparatively weak and also additionally masked by reduced adaptation.

### Including naturalistic features of odor stimuli does not increase non-synaptic interactions during short offset stimuli

Although our results up to now strongly indicate a limited usefulness of NSIs in processing temporal features of mixed odors, the stimuli are simple and do not share many features of natural odor plumes, which involve repeated exposures to odor filaments and fluctuations in concentration as well as whiff/blank duration. As a result of such stimulations, ORNs are likely to be in a state of partial adaptation under natural conditions, which may lead to NSIs becoming a more important feature. To investigate how such conditions affect NSIs, we performed two additional experiments with stimuli that more closely mimic the temporal structure and concentration fluctuations of natural plumes. In the first experiment, in order to simulate the repeated temporal features of a natural plume, ab3 ORNs were subjected to multiple simultaneous A and B odor pulses interspersed with brief clean air intervals **(Figure 9A and 9B)**. During this stimulation, the A ORN exhibited substantial adaptation, as reflected by a halving of its control firing rate. However, no corresponding increase in NSIs was observed, neither in absolute change in firing rate (*F*(2.999, 26.99) = 2.470, *p* = .08,*n* = 10, repeated measures ANOVA) nor in effective rate change (*F*(2.445, 22.00) = 0.4013, *p* = .7, repeated measures ANOVA). These findings suggest that, while repeated odor encounters produce notable adaptation, they do not enhance NSI strength.

**Figure 9.**
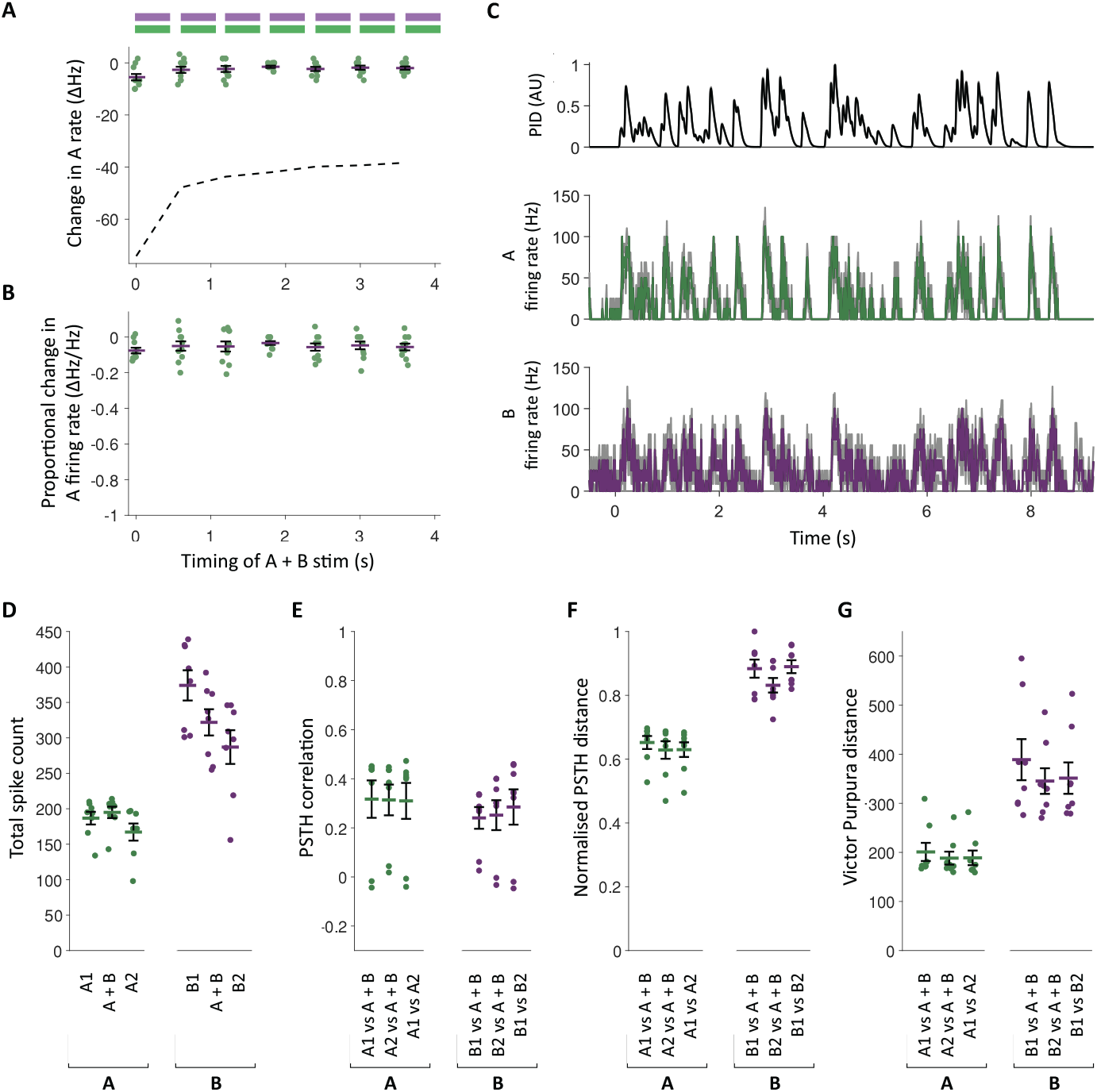
NSIs minimal during naturalistic odor stimuli. **A)** Response of ab3 ORNs to repeated paired stimuli, 500 ms B (purple bars) and A (green bars) pulse stimulations spaced 100 ms apart. Relationship between pulse offset from start of stimulation and change in A firing rate from control (no B pulse) to trial (with B pulse). Dashed line defines the maximum possible inhibition, defined as the negative mean control firing rate; *n* = 10. **B)** Relationship as panel A with change in A firing rate as a proportion of control firing rate. **C)** PSTH of mean responses of ab3 ORNs to single cognate odors during stimulation protocol A4: naturalistic plume with variable whiff duration and concentration. Plume features characterized by PID response to ethyl acetate (top); *n* = 8. Responses to protocol A4 during two controls (single cognate odor stimulation) and one trial (double odor stimulation) made by **D)** total spike counts; **E)** Pearson product-moment correlation between PSTHs; **F)** Euclidian distance between PSTHs, normalized by number of data points; and **G)** distance between spike trains, using the Victor-Purpura method. In each case, points represent individual A responses and error bars/shaded areas represent group mean ± SEM. A stimulation 10^−5^ and B stimulation 10^−4^.

To further investigate the potential impact of concentration fluctuations on NSIs, we conducted a second experiment simulating a naturalistic odor plume with variable concentrations and filament durations **(Figure 9C)**. A and B stimuli were delivered in this plume pattern, both individually and together, to mimic odor input from either a single source (fully correlated) or separate sources (uncorrelated). This approach offers our most realistic emulation of natural plume conditions and enables us to assess the potential role of NSIs in tasks such as odor source separation. The shorter simulation durations of this experimental protocol reduced ORN firing rates sufficiently to allow us to use an automated rather than manual spike sorting method. To assess both B-to-A and A-to-B inhibition, we utilized a spike sorting algorithm designed to recover masked spikes, enabling accurate measurement of activity in both ORNs (***Ellison et al., 2025***). As in the previous experiment, no significant differences were found between single and dual odor presentations; multiple analytical approaches consistently revealed no effect of odor pairing **(Figure 9D–9G)** (*p* > .2 for all analyses (Multiple One-way ANOVAs with Holm-Šídák correction). See **Table S1**) for test values

Simulated experiments replicating these more naturalistic odor stimulation conditions yielded similar results. During repeated paired stimuli, although the A response was substantially adapted by the repeated pulses, only a small degree of inhibition was observed **Figure 10A and 10B**. Peak firing rates were also slightly reduced during mixed naturalistic plumes **Figure 10C**. In both cases, NSI-mediated inhibition does not appear to increase during repeated odor encounters as these results are comparable to the unadapted conditions of short offset dual-odor stimulations (Figure 4). In order to rule out that these observations were due to the particular conditions of these stimulations, we repeated the paired odor pulse simulation with a range of pulse spacings from continuous to a gap sufficient for A to fully recover from adaptation before subsequent pulses. Although these simulations led to varying degrees of adaptation, the magnitude of NSIs remains largely unchanged **Figure 10G and 10H**. These results demonstrate that the nonlinear factors influencing NSIs are only likely to lead to high magnitude NSIs when only one of the paired ORNs has undergone sustained stimulation leading to an adapted state.

**Figure 10.**
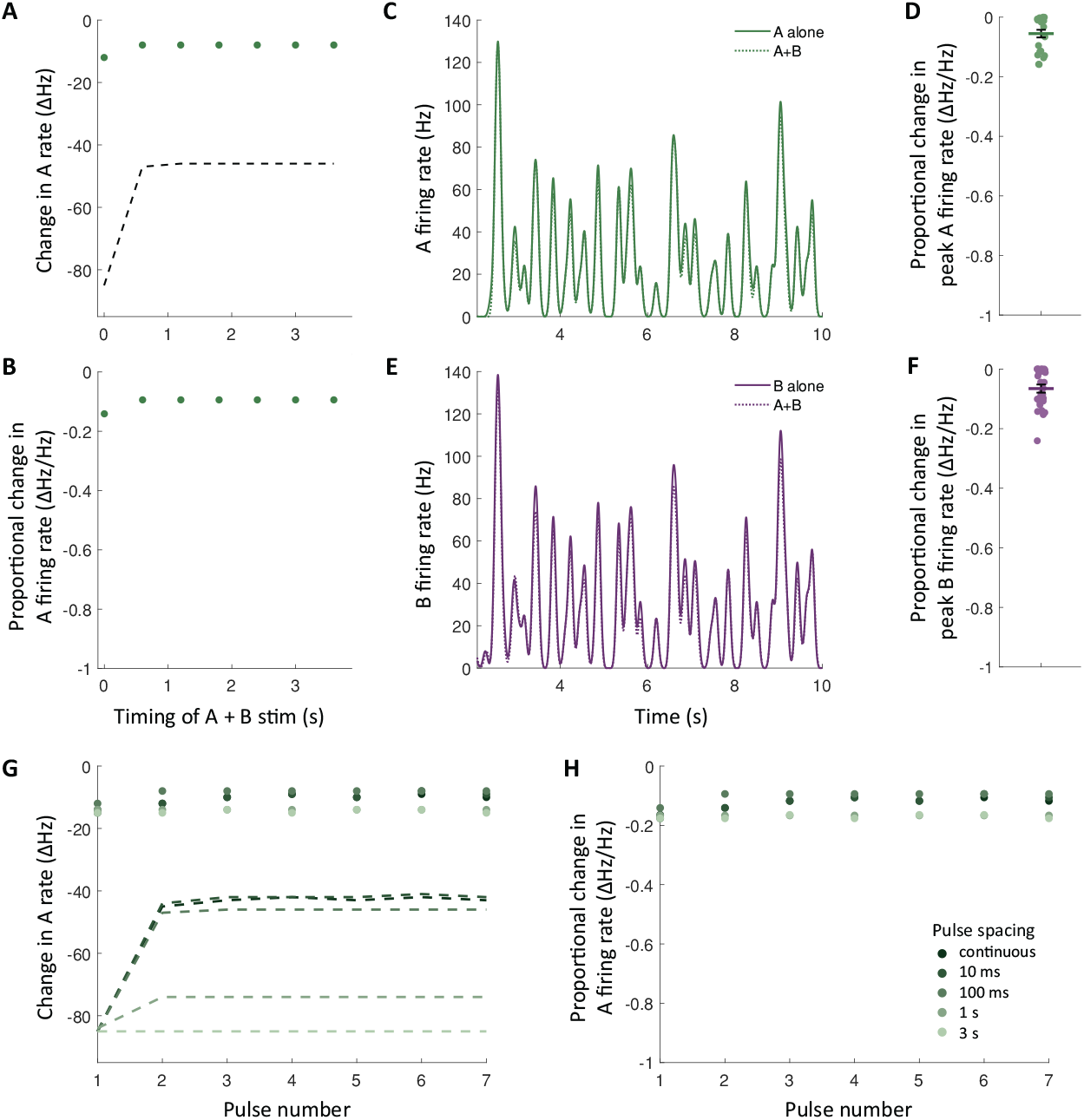
NSIs minimal during simulated naturalistic stimuli. **A)** Response of modeled ORNs to paired stimuli, same stimulation protocol Figure 9A. Relationship between pulse offset from start of stimulation and change in A firing rate from control (no B pulse) to trial (with B pulse). **B)** Relationship as panel A with change in A firing rate as a proportion of control firing rate. **C)** SDF of mean responses of modeled A ORN to single cognate odors during naturalistic plume simulation as Figure 9C. **D)** Relative peak firing response of A to simulated whiffs during A+B simulation. **E)** SDF of mean responses of modeled A ORN to single cognate odors during naturalistic plume simulation as Figure 9C. Responses to protocol A4 during two controls (single cognate odor stimulation) and one trial (double odor stimulation) made by **F)** Relative peak firing response of B to simulated whiffs during A+B simulation. **G)** Relationship between pulse offset from start of stimulation and change in A firing rate from control (no B pulse) to trial (with B pulse) at a range of pulse spacings from continuous (no space between pulses) to 3 s (when A fully recovers from adaptation between pulses). **H)** Relationship as panel G with change in A firing rate as a proportion of control firing rate. Dashed lines in A and G define mean maximum possible inhibition, defined as the negative mean control firing rate. In D and F, points represent individual peak responses and error bars/shaded areas represent group mean ± SEM.

In our analysis, we have examined changes in the A firing rate due to NSIs on the assumption that olfactory information is encoded primarily through rate coding. However, there is evidence for other forms of temporal coding in the insect olfactory system, such as first-spike latency, which is precise to the ms in insect ORNs and decreases with stimulation intensity (***Nagel and Wilson, 2011; Martelli et al., 2013; Egea-Weiss et al., 2018***). Therefore, as a final point, we return to two of our earlier experiments (**Figure 2** and **Figure 9A**) to examine the influence of NSIs on first spike latency. Overall, we find that NSIs increase first spike latency in a pattern consistent with our rate-based results **Figure S4**. During very short offset stimuli (A and B stimulations delivered almost simultaneously), first spike latency is largely unaffected by NSIs. As stimuli slow (with B delivered at a larger offset from the start of A), NSIs lead to an increase in first spike latency, consistent with a larger proportional reduction in the A firing rate. During repeated paired stimuli, where odors stimulating both ORNs are delivered simultaneously, although first spike latency increases over time, the increase happens equally in control conditions and when odor B is delivered simultaneously. This is consistent with the minimal changes of the A firing rate during simultaneous A and B stimulations compared to A stimulation alone.

## Discussion

In this paper, we reported the results of an in-depth investigation of the phenomenology of NSIs in the olfactory sensilla of *Drosophila*. We focused our observations on the spiking activity of ORNs as measured by SSRs. This spiking activity is presumably the only information that is transmitted outside the sensillum and into the brain. We found that NSIs measured as ORN firing changes occur whenever the ORNs in a sensillum are both active at the same time. However, the effective magnitude of the interaction, as measured in the proportional change of the number of fired spikes, is highly dependent on the history of ORN activation. In essence, significant NSIs are only observed when one of the ORNs has adapted due to sustained or repeated activation by odorants and the other ORN has not adapted. In this situation the second ORN can significantly reduce the firing of the first. In all other situations, NSIs are quite weak.

These findings are consistent with ephaptic interactions observed in other insect sensory modalities. In paired gustatory receptor neurons (GRNs), which share a similar sensillar housing arrangement to ORNs, inhibition also appears to depend on adaptation, with large proportional reductions of the GRN firing rate only occurring following adaptation (***Lee et al., 2025***). Similarly, ephaptic lateral inhibition among achromatic fly photoreceptors operates over relatively slow timescales, acting as a high-pass filter and suppressing redundant low-frequency signals (***Weckström and Laughlin, 2010***). Despite the demand for rapid processing in vision, here too these interactions appear to modulate longer-timescale signals that parallel our findings in the olfactory system.

In order to understand the extent to which electrical interactions alone can explain the adaptation-dependence of NSIs, we built a detailed conductance-based model of two ORNs co-housed in a sensillum. We were able to reproduce our experimental observations and subsequently dissect the mechanism underlying this phenomenology. This analysis revealed that the most likely explanation for our observations is that the most noticeable NSIs occur for an adapted inhibited neuron and non-adapted inhibiting neuron because a high firing rate response in the inhibiting neuron causes a significant drop in sensillum LFP and the inhibited neuron is particularly sensitive to this because it has wide open receptors and hence a large change in receptor current while at the same time firing at lower rate due to adaptation. At this lower rate, the F-I curve is steep so that the change of receptor current has strong effects on the resulting firing rate. In other situations, either the change in receptor current is small (low odor concentrations) or the F-I curve is flat (the non-adapted ORN is responding to high concentrations with its maximal firing rate). In either case, there is little change in ORN firing rate due to NSIs. Interestingly, when stimulations are concurrent, the model shows very little effect; similar ideas were also discussed by ***Su et al. (2011***), but with respect to a different mechanism of spike response depending on the rate of change of the membrane potential. Our model describes LFP responses indicative of NSIs: LFP deflections are smaller during concurrent stimulation of both ORNs in comparison to the linear sums of responses during single stimulations **Figure S5**, similar to experiments (***Zhang et al., 2019***). However, our work demonstrates that NSIs cannot be predicted from the LFP alone, as has previously been assumed. Instead, we argue that changes in spike frequency adaptation current during periods of rapid adaptation oppose these effects.

Our observations have implications for the putative role of NSIs in olfactory information processing. Because neurons are grouped systematically in sensilla it is likely that NSIs play a functional role selected through evolution. Several such roles have been suggested in the literature, most prominently extending dynamic range (***Vermeulen and Rospars, 2004***), novelty detection (***Todd and Baker, 1999; Su et al., 2012***), interpreting countervailing cues (***Wu et al., 2022; Puri et al., 2024***) and pre-processing of rapidly changing olfactory mixture stimuli in turbulent plumes (***Stierle et al., 2013; Bandyopadhyay and Sachse, 2023; Raiser et al., 2024***). Our results do not directly speak to the dynamic range hypothesis, though previous modeling has cast some doubts (***Pannunzi and Nowotny, 2021***). However, the observation of weak NSIs for short offset stimuli, which persists even if adaptation comes into play through repeated ORN activation in a plume, does not support the idea of NSIs playing a major role in plume mixture processing. It is much more likely that they are mainly used in novelty detection.

We have interpreted our findings with the help of a conductance-based model of two ORNs co-housed in a sensillum, assuming a generic M-current-based spike rate adaptation mechanism. The description for the sodium and delayed rectifier current was taken from a hippocampal pyramidal neuron model (***Traub and Miles, 1991***) as a generic, type 1, numerically well-behaved spiking neuron model. While these details almost certainly do not exactly match the *Drosophila* electrophysiology, we found that the model matched all of our experimental observations with minimal initial tuning. This gives us confidence that the model captures the relevant electrical properties well enough to be a useful tool for interpretation. The mechanistic explanation of our observations resulting from the analysis of the model furthermore does make intuitive sense beyond the details of the model further supporting our interpretation.

Our findings appear to contrast with the conclusions of a recent study examining the same question of the effect of NSIs during short offset stimuli (***Raiser et al., 2024***), which reported appreciable inhibition of the B neuron by A stimulation but not vice versa. These measurements were made for small onset delays (0-96 ms). Here, we have refrained from reporting experimental results concerning the effects of A activity on the B firing rate because we judged that spike sorting the smaller B spikes during stimulation of both neurons is too unreliable, and missing B spikes during spike sorting can be misinterpreted as B being inhibited. With respect to the effect of B stimulation on the activity of A, both studies agree that NSIs have little effect on A firing rate for short onset delays. Our computational model suggests that inhibition should affect both neurons similarly, though there can be quantitative differences due to neurons having different sizes and hence differently strong influence over the LFP.

In conclusion, we have demonstrated in experiments and with a computational model that NSIs have little effect on ORN firing for short odor onset delays and conclude that it is likely that NSIs play less of a role in the processing of turbulent plumes and more of a role in novelty detection.

## Acknowledgments

We thank Claudio Alonso for his help with fly experiments and for providing lab support. The work in this paper was funded by a Leverhulme Trust project grant (TN and GK) and the EPSRC (Brains on Board project, Grant Number EP/P006094/1, ActiveAI project, Grant Number EP/S030964/1) (TN).

## Author contributions

**Lydia Ellison**: Conceptualization, Investigation, Methodology, Visualization, Writing - original draft. **György Kemenes**: Funding acquisition, Conceptualization, Supervision, Writing - review and editing. **Thomas Nowotny**: Conceptualization, Funding acquisition, Methodology, Software, Supervision, Writing - original draft.

## Ethical approval

The research reported in this manuscript has received ethics approval from the ANIMAL WELFARE AND ETHICAL REVIEW BODY (AWERB) of the University of Sussex.

## Supplementary Information

**Figure S1.**
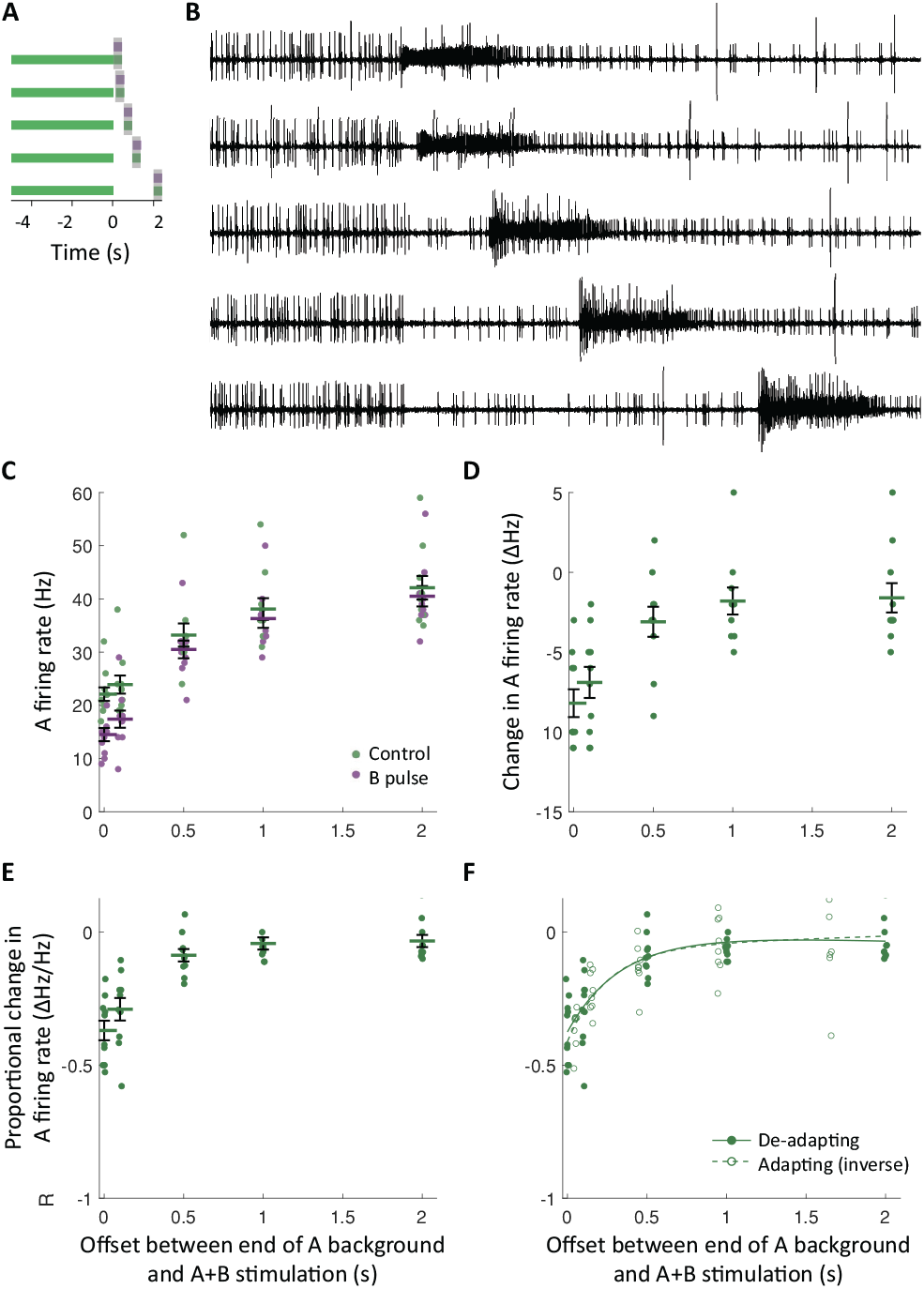
De-adaption recapitulates adaptation dependence of NSIs. **A)** De-adapting stimulation protocol: 500 ms paired A and B pulses (small purple bars in grey sampling window) offset from end of 5 s adapting A stimulations (long green bars). **B)** Example SSR data illustrating series of stimulations with increasing offset of paired pulses from end of adapting A stimulations. **C)** Comparison of ab3A response to De-adapting protocol: A stimulation alone (control) and with B pulse, *n* = 10. Here and in the following panels points represent individual ab3A responses and error bars represent group mean ± SEM. **D)** Relationship between offset of paired pulses from end of A stimulation and change in ab3A firing rate from control (no B pulse) to trial (with B pulse). Data from panel C. **E)** Relationship as panel D, with ab3A inhibition as proportional change in firing rate. **F)** Comparison of ab3A response to De-adapting protocol (filled points and solid line) and Adapting stimulation protocol (hollow points and dashed line): first 5 offsets of data from Figure 2F inverted and aligned by mean. Lines define two-phase exponential models fit to each data set.

**Figure S2.**
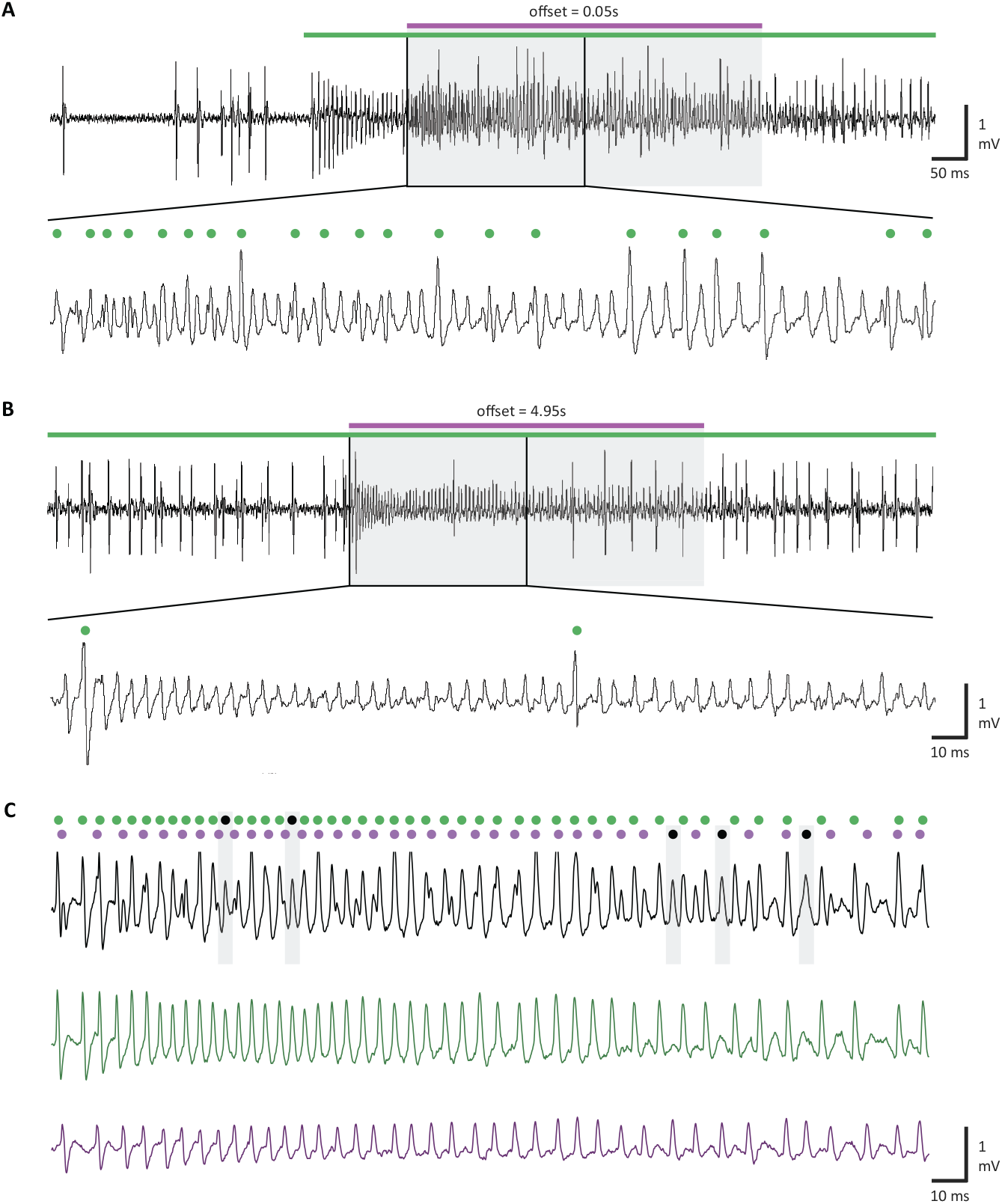
Simulated potentials in the olfactory sensillum model. When stimulation of ab3B begins soon (50 ms) after the start ab3A stimulation, spike sorting is more challenging due to a large number of ambiguous spike shapes **(A)**. Green circles indicated A spikes identified by manual sorting. This is even more pronounced during concurrent stimulation (not shown). When the offset between the stimulations is longer (4.95 s) and ab3A is in an adapted low firing regime, A spikes are clearly identifiable **(B)**. Generation of ‘synthetic’ SSR data from a two neuron sensillum can be achieved by combining traces from sensilla where only one ORN is active **(C)**. Composite data (black trace) formed from the linear addition of individual traces from ab3 sensilla with ab3A inactive (purple = ab3B only) and with ab3B inactive (green = ab3A only). Such composite data with corresponding ground truth spike identification (green and purple circles) illustrates that during high firing rates, ambiguous spike shapes can lead to errors in ‘A’ spike sorting both by reducing the apparent size of A spikes such that they are likely to be rejected as ‘B’ and by increasing the size of B spikes such that they are likely to be mistaken for ‘A’ spikes. Likely errors indicated by black circles and corresponding shaded boxes over relevant spikes. SSR data was collected from heterozygous fly lines expressing the inward rectifier potassium channel Kir2.1 in single ORNs of the ab3 sensillum: w*;UASKCNJ2;Or22a-GAL4 and w*;Or85b-GAL4;UAS-KCNJ2. Expression of this channel results in hyperpolarisation and inactivity, thus leaving ab3 sensilla with only one active ORN ***Retzke et al. (2017***).

**Figure S3.**
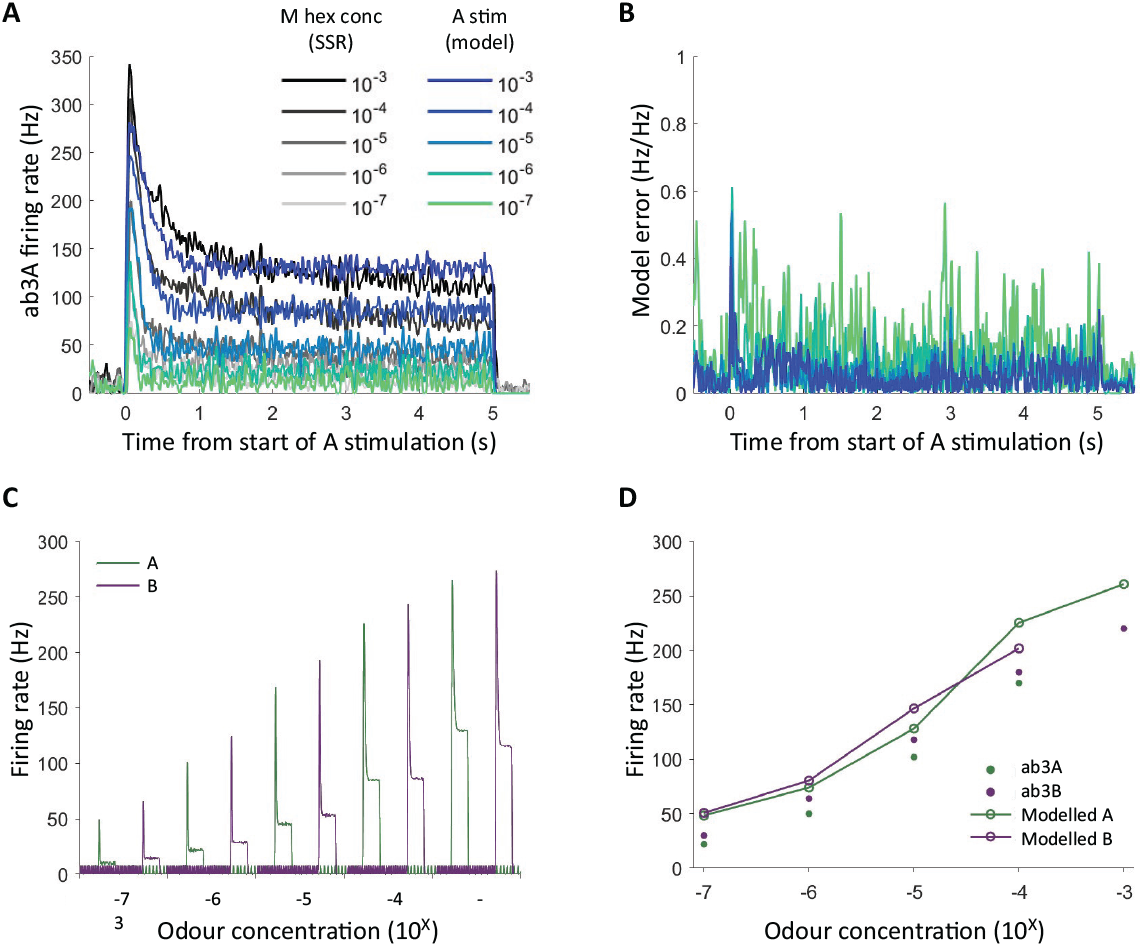
Evaluation of model fit to ab3 ORN responses. **A)** Comparison of simulated A and actual ab3A responses to a range of odor dilutions using parameters determined by fitting with *scipy*.*optimize*.*minimize*. **B)** Normalized error between simulated A and actual ab3A responses shown in (A). **C)** Responses of simulated A and B to 3 s simulations at a range of odor concentrations. Parameters used as in simulations aiming to replicate experimental conditions, with asymmetry reflecting actual morphological differences and B parameters manually fit to actual ab3B responses. **D)** Comparison between dose responses of simulated A and B and actual ab3 ORNs with same parameters as (C). Responses averaged over 500 ms sampling window.

**Figure S4.**
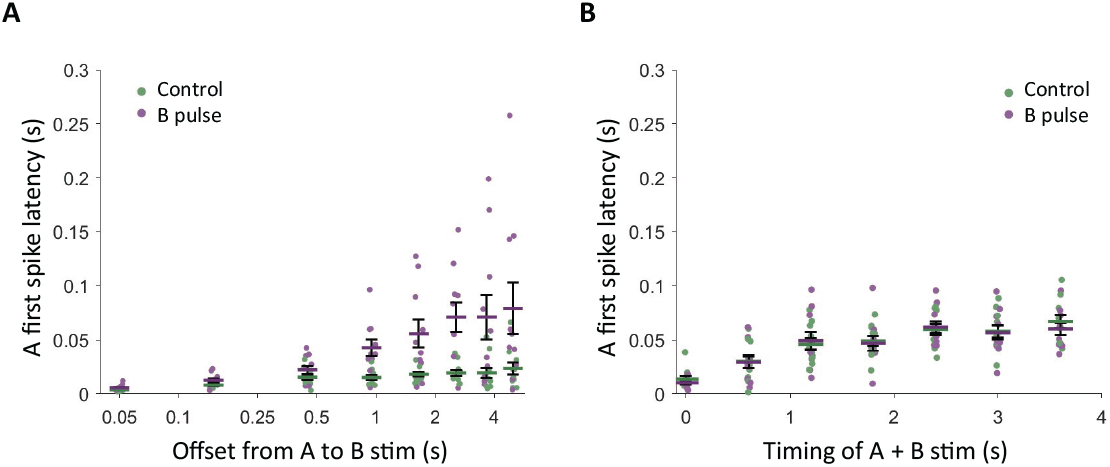
First spike latency reduced by NSIs. **A)** First spike latency during short offset stimuli (Figure 2) control (no B pulse) and trial (with B pulse) of short offset stimulation protocol in ab3. **B)** First spike latency during repeated odor encounters (Figure 9A) control (no B pulse) and trial (with B pulse) of repeated paired stimuli.

**Figure S5.**
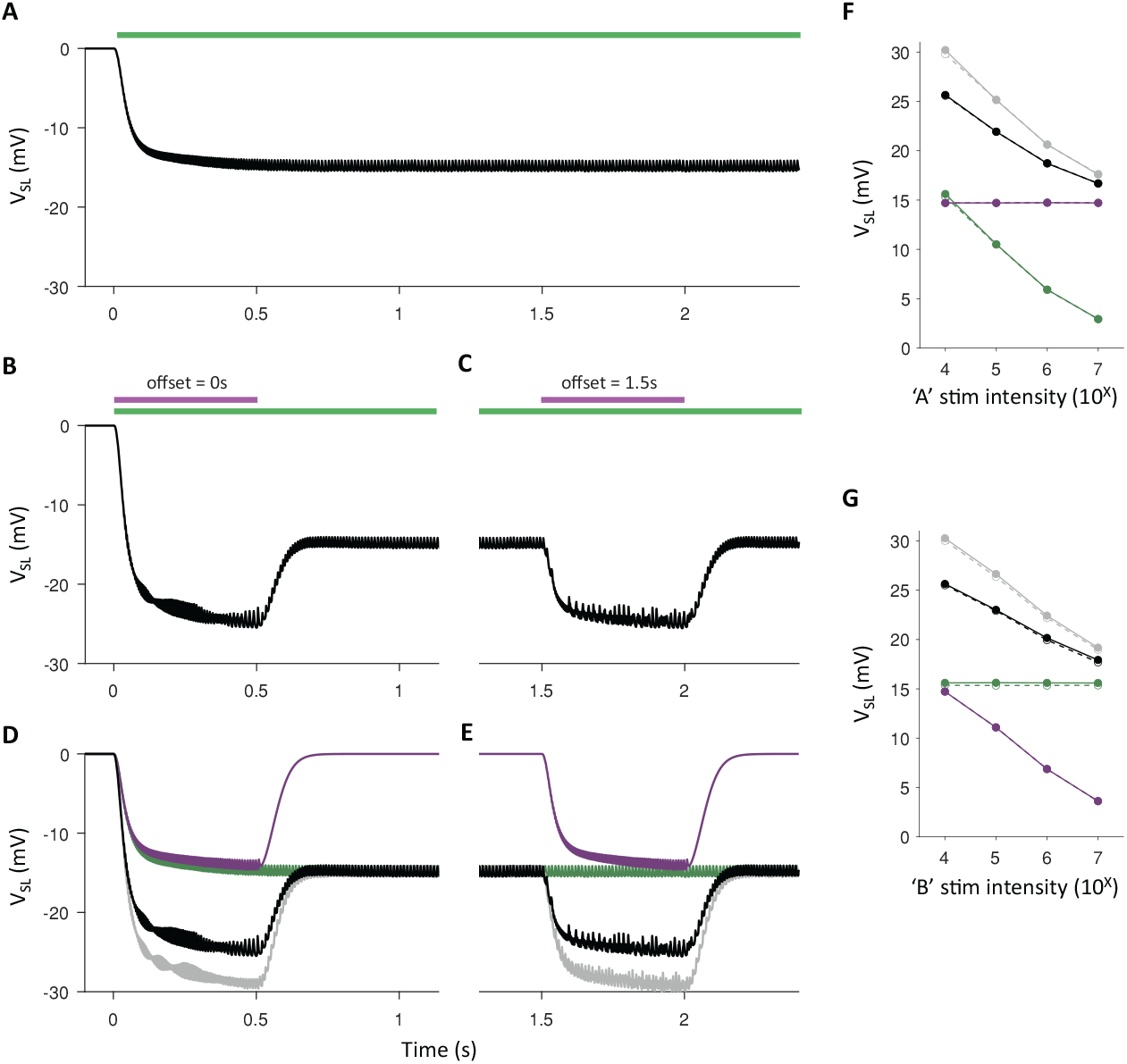
Magnitude of LFP deflection independent of stimulus timing. **A)** Time-resolved simulation of LFP (*V*_SL_) during A stimulation alone (green bar). **B)** As panel A during experimental trial with concurrent stimulation of A and B (purple bar). **C)** As panel B following long offset before B stimulation. Note that the LFP deflections in panels B and C are of a similar magnitude. **D)** LFP during A stimulation alone (green); B stimulation alone (purple; A and B stimulation (black); and prediction based on linear addition of responses to individual A and B stimulations (gray). As previously demonstrated in ***Zhang et al. (2019***), NSIs lead to a smaller LFP deflection than predicted during concurrent A and B stimulation. **E)** As panel D following long offset before B stimulation. Note that the discrepancy between predicted and simulated LFP deflections in panels D and E are comparable. **F)** Peak LFP responses (absolute values) plotted as a function of A stimulation intensity. Data colored as panels D & E. Hollow points and dashed line correspond to concurrent stimulation of A and B (as panel D). Filled points and solid line correspond to long offset B stimulation (as panel E). **G)** As panel F, responses as a function of B stimulation intensity. Note that in both panels F and G, concurrent and long offset stimulations lead to near identical LFP responses.

**Table S1.**
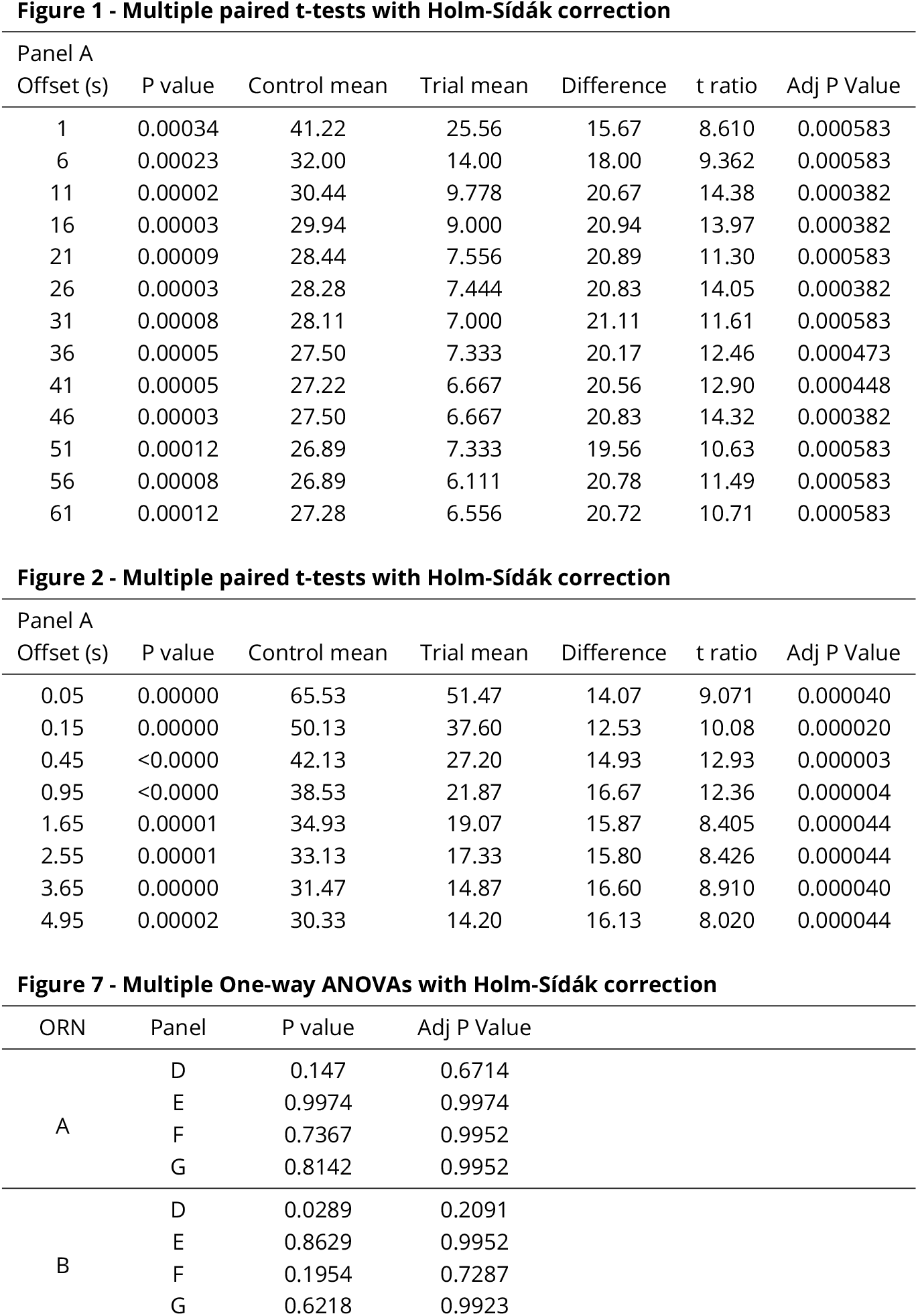
Summary statistics.

1 - Multiple paired t-tests with Holm-Sídák correction
| Panel A |  |  |  |  |  |  |
| --- | --- | --- | --- | --- | --- | --- |
| Offset (s) | P value | Control mean | Trial mean | Difference | t ratio | Adj P Value |
| 1 | 0.00034 | 41.22 | 25.56 | 15.67 | 8.610 | 0.000583 |
| 6 | 0.00023 | 32.00 | 14.00 | 18.00 | 9.362 | 0.000583 |
| 11 | 0.00002 | 30.44 | 9.778 | 20.67 | 14.38 | 0.000382 |
| 16 | 0.00003 | 29.94 | 9.000 | 20.94 | 13.97 | 0.000382 |
| 21 | 0.00009 | 28.44 | 7.556 | 20.89 | 11.30 | 0.000583 |
| 26 | 0.00003 | 28.28 | 7.444 | 20.83 | 14.05 | 0.000382 |
| 31 | 0.00008 | 28.11 | 7.000 | 21.11 | 11.61 | 0.000583 |
| 36 | 0.00005 | 27.50 | 7.333 | 20.17 | 12.46 | 0.000473 |
| 41 | 0.00005 | 27.22 | 6.667 | 20.56 | 12.90 | 0.000448 |
| 46 | 0.00003 | 27.50 | 6.667 | 20.83 | 14.32 | 0.000382 |
| 51 | 0.00012 | 26.89 | 7.333 | 19.56 | 10.63 | 0.000583 |
| 56 | 0.00008 | 26.89 | 6.111 | 20.78 | 11.49 | 0.000583 |
| 61 | 0.00012 | 27.28 | 6.556 | 20.72 | 10.71 | 0.000583 |

**Figure 2 - Multiple paired t-tests with Holm-Sídák correction**
| Panel A |  |  |  |  |  |  |
| --- | --- | --- | --- | --- | --- | --- |
| Offset (s) | P value | Control mean | Trial mean | Difference | t ratio | Adj P Value |
| 0.05 | 0.00000 | 65.53 | 51.47 | 14.07 | 9.071 | 0.000040 |
| 0.15 | 0.00000 | 50.13 | 37.60 | 12.53 | 10.08 | 0.000020 |
| 0.45 | <0.0000 | 42.13 | 27.20 | 14.93 | 12.93 | 0.000003 |
| 0.95 | <0.0000 | 38.53 | 21.87 | 16.67 | 12.36 | 0.000004 |
| 1.65 | 0.00001 | 34.93 | 19.07 | 15.87 | 8.405 | 0.000044 |
| 2.55 | 0.00001 | 33.13 | 17.33 | 15.80 | 8.426 | 0.000044 |
| 3.65 | 0.00000 | 31.47 | 14.87 | 16.60 | 8.910 | 0.000040 |
| 4.95 | 0.00002 | 30.33 | 14.20 | 16.13 | 8.020 | 0.000044 |

**Figure 7 - Multiple One-way ANOVAs with Holm-Sídák correction**
| ORN | Panel | P value | Adj P Value |
| --- | --- | --- | --- |
| A | D | 0.147 | 0.6714 |
|  | E | 0.9974 | 0.9974 |
|  | F | 0.7367 | 0.9952 |
|  | G | 0.8142 | 0.9952 |
| B | D | 0.0289 | 0.2091 |
|  | E | 0.8629 | 0.9952 |
|  | F | 0.1954 | 0.7287 |
|  | G | 0.6218 | 0.9923 |

**Table S2.** Model parameters.

| ORN parameters |  |
| --- | --- |
| $E_K$ | −95 mV |
| $E_{Na}$ | 50 mV |
| $E_{leak}$ | −63.563 mV |
| * $C$ | 1 nF |
| * $C_D$ | 1 nF |
| * $g_{leak}$ | 0.186 85 μS |
| * $g_{Na}$ | 50 μS |
| * $g_K$ | 10 μS |
| * $g_{VV}$ | 1 μS |
| # $I_{base}$ | 0.8 nA |
| Receptor current parameters |  |
| $^{\dagger} cmid$ | −3.34225086 |
| $^{\dagger} cslope$ | 0.877785161 |
| # $cmid_B$ | $cmid * 1.2$ |
| # $cslope_B$ | $cslope$ |
| * $^{\dagger} g_R$ | 0.975 556 066 μS |
| $\tau_{SR}$ | 25 ms |
| M-current parameters |  |
| $^{\dagger} P_{M1}$ | $4.228\ 252\ 45 \times 10^{-2}$ |
| $^{\dagger} P_{M2}$ | 21.540 256 2 mV |
| $^{\dagger} P_{M3}$ | 7.988 890 05 mV |
| $^{\dagger} P_{M4}$ | $1.164\ 135\ 90 \times 10^{-3}$ |
| $^{\dagger} P_{M5}$ | −31.591 666 2 mV |
| $^{\dagger} P_{M6}$ | −4.515 283 81 mV |
| * $^{\dagger} g_M$ | 0.844 657 433 μS |
| Sensillum parameters |  |
| $g_{SL, leak}$ | 1 μS |
| $E_{SL, leak}$ | −30 mV |
| $C_{SL, leak}$ | 8 nF |

$^{\dagger}$ fit to ab3A responses to methyl hexanoate using *scipy.optimize.minimize*.
\* scaled by 0.74 to account for ab3B morphology.

## References

Andersson MN, Larsson MC, Blaženec M, Jakuš R, Zhang QH, Schlyter F. Peripheral modulation of pheromone response by inhibitory host compound in a beetle. Journal of Experimental Biology. 2010 oct; 213(19):3332–3339. doi: 10.1242/jeb.044396.

Bandyopadhyay P, Sachse S. Mixing things up! — how odor blends are processed in Drosophila. Current Opinion in Insect Science. 2023 oct; 59:101099. doi: 10.1016/j.cois.2023.101099.

Benton R, Dahanukar A. Electrophysiological recording from Drosophila olfactory Sensilla. Cold Spring Harbor Protocols. 2011 jul; 6(7):824–838. doi: 10.1101/pdb.prot5630.

Cafaro J. Multiple sites of adaptation lead to contrast encoding in the Drosophila olfactory system. Physiological Reports. 2016 apr; 4(7):1–14. doi: 10.14814/phy2.12762.

Celani A, Villermaux E, Vergassola M. Odor landscapes in turbulent environments. Physical Review X. 2014; 4(4). doi: 10.1103/PhysRevX.4.041015.

Chang H, Guo M, Wang B, Liu Y, Dong S, Wang G. Sensillar expression and responses of olfactory receptors reveal different peripheral coding in two Helicoverpa species using the same pheromone components. Scientific Reports. 2016; 6(July 2015):1–12. doi: 10.1038/srep18742.

Cossé AA, Todd JL, Baker TC. Neurons discovered in male Helicoverpa zea antennae that correlate with pheromone-mediated attraction and interspecific antagonism. Journal of Comparative Physiology - A Sensory, Neural, and Behavioral Physiology. 1998; 182(5):585–594. doi: 10.1007/s003590050205.

Egea-Weiss A, Renner A, Kleineidam CJ, Szyszka P. High Precision of Spike Timing across Olfactory Receptor Neurons Allows Rapid Odor Coding in Drosophila. iScience. 2018 jun; 4:76–83. doi: 10.1016/j.isci.2018.05.009.

Ellison L, Raiser G, Garrido-Peña A, Kemenes G, Nowotny T. SSSort 2.0: A semi-automated spike detection and sorting system for single sensillum recordings. Journal of Neuroscience Methods. 2025; 415(July 2024):110351. doi: 10.1016/j.jneumeth.2024.110351.

Faber DS, Korn H. Electrical field effects: Their relevance in central neural networks. Physiological Reviews. 1989; 69(3):821–863. doi: 10.1152/physrev.1989.69.3.821.

Gomez-Diaz C, Martin F, Garcia-Fernandez JM, Alcorta E. The two main olfactory receptor families in drosophila, ORs and IRs: A comparative approach. Frontiers in Cellular Neuroscience. 2018 aug; 12:253. doi: 10.3389/fncel.2018.00253.

Hallem EA, Carlson JR. Coding of Odors by a Receptor Repertoire. Cell. 2006; 125(1):143–160. doi: 10.1016/j.cell.2006.01.050.

Jefferys JGR. Nonsynaptic modulation of neuronal activity in the brain: Electric currents and extracellular ions. Physiological Reviews. 1995; 75(4):689–723. doi: 10.1152/physrev.1995.75.4.689.

Jordán MJ, Tandon K, Shaw PE, Goodner KL. Aromatic profile of aqueous banana essence and banana fruit by gas chromatography-mass spectrometry (GC-MS) and gas chromatography-olfactometry (GC-O). Journal of Agricultural and Food Chemistry. 2001; 49(10):4813–4817. doi: 10.1021/jf010471k.

Kaissling KE. Chemo-electrical transduction in insect olfactory receptors. Annual Review of Neuroscience. 1986; VOL. 9:121–145. doi: 10.1146/annurev.neuro.9.1.121.

Lee MH, Park SH, Joo KM, Kwon JY, Lee KH, Kang KJ. Drosophila hcn mediates gustatory homeostasis by preserving sensillar transepithelial potential in sweet environments. eLife. 2024; 13:1–19. doi: 10.7554/eLife.96602.

Lee M, Kima SY, Parka T, Yoone SE, Kime YJ, Joob KM, Kwonc JY, Kimd K, Kanga KJ. An evolutionarily conserved cation channel tunes the sensitivity of gustatory neurons to ephaptic inhibition in Drosophila. Proceedings of the National Academy of Sciences. 2025;122(3). doi: 10.1073/pnas.2413134122.

Martelli C, Carlson JR, Emonet T. Intensity invariant dynamics and odor-specific latencies in olfactory receptor neuron response. Journal of Neuroscience. 2013 apr; 33(15):6285–6297. doi: 10.1523/JNEUROSCI.0426-12.2013.

Martelli C, Fiala A. Slow presynaptic mechanisms that mediate adaptation in the olfactory pathway of Drosophila. eLife. 2019 jun; 8. doi: 10.7554/eLife.43735.

Martin F, Alcorta E. Measuring activity in olfactory receptor neurons in Drosophila: Focus on spike amplitude. Journal of Insect Physiology. 2016 dec; 95:23–41. doi: 10.1016/j.jinsphys.2016.09.003.

Nagel KI, Wilson RI. Biophysical mechanisms underlying olfactory receptor neuron dynamics. Nature Neuroscience. 2011 feb; 14(2):208–218. doi: 10.1038/nn.2725.

Ng R, Wu ST, Su CY. Neuronal Compartmentalization: A Means to Integrate Sensory Input at the Earliest Stage of Information Processing? BioEssays. 2020; 42(8). doi: 10.1002/bies.202000026.

Olsson SB, Hansson BS. Electroantennogram and single sensillum recording in insect antennae. Methods in Molecular Biology. 2013; 1068:157–177. doi: 10.1007/978-1-62703-619-1_11.

Pannunzi M, Nowotny T. Non-synaptic interactions between olfactory receptor neurons, a possible key feature of odor processing in flies. PLoS Computational Biology. 2021 jun; 17(12):e1009583. doi: 10.1371/journal.pcbi.1009583.

Pellegrino M, Nakagawa T, Vosshall LB. Single sensillum recordings in the insects Drosophila melanogaster and Anopheles gambiae. Journal of Visualized Experiments. 2010 feb; (36). doi: 10.3791/1725.

Puri P, Wu ST, Su CY, Aljadeff J. Peripheral preprocessing in Drosophila facilitates odor classification. Proceedings of the National Academy of Sciences of the United States of America. 2024; 121(21):1–11. doi: 10.1073/pnas.2316799121.

Raiser G, Galizia CG, Szyszka P. A high-bandwidth dual-channel olfactory stimulator for studying temporal sensitivity of olfactory processing. Chemical Senses. 2017; 42(2):141–151. doi: 10.1093/chemse/bjw114.

Raiser G, Galizia CG, Szyszka P. Olfactory receptor neurons are sensitive to stimulus onset asynchrony: implications for odor source discrimination. Chemical Senses. 2024 08; 49:bjae030. doi: 10.1093/chemse/bjae030.

Retzke T, Thoma M, Hansson BS, Knaden M. Potencies of effector genes in silencing odor-guided behavior in Drosophila melanogaster. Journal of Experimental Biology. 2017 may; 220(10):1812–1819. doi: 10.1242/jeb.156232.

Schmidt HR, Benton R. Molecular mechanisms of olfactory detection in insects: Beyond receptors: Insect olfactory detection mechanisms. Open Biology. 2020; 10(10). doi: 10.1098/rsob.200252rsob200252.

Shaw SR. Retinal resistance barriers and electrical lateral inhibition. Nature. 1975; 255(5508):480–483. doi: 10.1038/255480a0.

Stierle JS, Giovanni Galizia C, Szyszka P. Millisecond stimulus onset-asynchrony enhances information about components in an odor mixture. Journal of Neuroscience. 2013; 33(14):6060–6069. doi: 10.1523/JNEUROSCI.5838-12.2013.

Stimberg M, Brette R, Goodman DF. Brian 2, an intuitive and efficient neural simulator. eLife. 2019 aug; 8:e47314. doi: 10.7554/eLife.47314.

Su CY, Martelli C, Emonet T, Carlson JR. Temporal coding of odor mixtures in an olfactory receptor neuron. Proceedings of the National Academy of Sciences. 2011 mar; 108(12):5075–5080. doi: 10.1073/PNAS.1100369108.

Su CY, Menuz K, Reisert J, Carlson JR. Non-synaptic inhibition between grouped neurons in an olfactory circuit. Nature. 2012 ec; 492(7427):66–71. doi: 10.1038/nature11712.

Takeichi Y, Uebi T, Miyazaki N, Murata K, Yasuyama K, Inoue K, Suzaki T, Kubo H, Kajimura N, Takano J, Omori T, Yoshimura R, Endo Y, Hojo MK, Takaya E, Kurihara S, Tatsuta K, Ozaki K, Ozaki M. Putative Neural Network Within an Olfactory Sensory Unit for Nestmate and Non-nestmate Discrimination in the Japanese Carpenter Ant: The Ultra-structures and Mathematical Simulation. Frontiers in Cellular Neuroscience. 2018 sep; 12:310. doi: 10.3389/fncel.2018.00310.

Todd JL, Baker TC. Function of Peripheral Olfactory Organs. In: Hansson BS, editor. Insect Olfaction Berlin, Heidelberg: Springer Berlin Heidelberg; 1999. p. 67–96. doi: 10.1007/978-3-662-07911-9_4.

Traub RD, Miles R. Neuronal Networks of the Hippocampus. Cambridge University Press; 1991.

Vermeulen A, Rospars JP. Why are insect olfactory receptor neurons grouped into sensilla? The teachings of a model investigating the effects of the electrical interaction between neurons on the transepithelial potential and the neuronal transmembrane potential. European Biophysics Journal. 2004 nov; 33(7):633–643. doi: 10.1007/s00249-004-0405-4.

Weckström M, Laughlin S. Extracellular potentials modify the transfer of information at photoreceptor output synapses in the blowfly compound eye. Journal of Neuroscience. 2010; 30(28):9557–9566. doi: 10.1523/JNEUROSCI.6122-09.2010.

Wu ST, Chen JY, Martin V, Ng R, Zhang Y, Grover D, Greenspan RJ, Aljadeff J, Su CY. Valence opponency in peripheral olfactory processing. Proceedings of the National Academy of Sciences of the United States of America. 2022 feb; 119(5). doi: 10.1073/pnas.2120134119.

Xu P, Choo YM, Chen Z, Zeng F, Tan K, Chen TY, Cornel AJ, Liu N, Leal WS. Odorant Inhibition in Mosquito Olfaction. iScience. 2019 sep; 19:25–38. doi: 10.1016/j.isci.2019.07.008.

Yee E, Chan R, Kosteniuk PR, Chandler GM, Biltoft CA, Bowers JF. Measurements of level-crossing statistics of concentration fluctuations in plumes dispersing in the atmospheric surface layer. Boundary-Layer Meteorology. 1995; 73(1-2):53–90. doi: 10.1007/BF00708930.

Zhang Y, Tsang TK, Bushong EA, Chu LA, Chiang AS, Ellisman MH, Reingruber J, Su CY. Asymmetric ephaptic inhibition between compartmentalized olfactory receptor neurons. Nature Communications. 2019 dec; 10(1). doi: 10.1038/s41467-019-09346-z.

